# Perceived social value is more strongly linked to gossip decisions with age in adolescence

**DOI:** 10.64898/2026.10.10.757213

**Authors:** Seungyoon Julie Oh, Youngjo Song, Jeungmin Lee, Yueun Jung, Jaeseung Jeong

## Abstract

Gossip is central to social exchange, yet how adolescents select what to share and with whom remains unclear. Across two endent samples aged 10–18 years (behavioural, n = 98; fMRI, n = 101 scanned), adolescents read information about absent third parties, chose whether and with whom to share it, and rated interest, usefulness and emotional response. Sharing frequency showed no detectable age association. The association between interest ratings and sharing strengthened with age in both samples; usefulness showed a smaller age interaction. Adolescents selected close friends more often with age when sharing. During scenario reading, before participants reported choices, BOLD responses in a left inferior frontal cluster tracked subsequent interest ratings more strongly with age (n = 92), also within regions defined by a mental-state judgment localizer. Together, these findings link age in adolescence to stronger valuation-to-decision coupling for interest and a greater proportion of sharing directed to close friends.

---

Social information becomes available to others through choices about what to pass on and whom to tell. Such choices help determine who can learn about whom, making information sharing an important component of social interaction. Gossip, the transmission of information about absent third parties, provides a tractable setting for studying these decisions^1,2^. Through gossip, people exchange reputational information, learn about others and participate in relationships^3,4^. Examining adolescents as providers of this information adds a complementary perspective to research on how they respond to and learn from their social environments^5,6,7^.

Children share information about absent others^8^ and begin to understand how gossip contributes to reputations^9,10^. During adolescence, peer relationships and reputational concerns become increasingly prominent^11,12^. Research on information sharing shows that people consider the usefulness and emotional consequences of information, as well as the receiver’s uncertainty^13^. Recent work also demonstrates that knowledge of social network structure informs strategic gossip^14^. An unresolved developmental question is how adolescents’ appraisals are associated with the information they choose to share, alongside their choices of recipients. Sharing frequency alone cannot answer this question: individuals who share equally often may differ in which information they select and whom they choose to tell.

We distinguish three reported appraisals of social information: interest, usefulness and emotional response. Value-based decision frameworks and studies of information seeking motivate examining these appraisals separately, because information can have both instrumental and non-instrumental value^15,16,17^. We quantify the within-participant association between these appraisals and sharing choices, which we refer to as valuation-to-decision coupling. This approach allows us to examine whether age is associated with stronger correspondence between appraisals and sharing choices overall, and whether interest makes a greater relative contribution. We also examine recipient selection as a distinct aspect of sharing. Because appraising information for communication may involve understanding other people^18,19,20^, we assess whether valuation-to-decision coupling for interest covaries with mental-state inference performance.

We studied these questions in two independent cohorts aged 10–18 years using personalised scenarios about absent third parties, adapting an established gossip-sharing task^21^. Participants chose whether to share each scenario, selected a recipient when sharing, and subsequently rated their appraisals. We combined models of individual trial choices with validation across participants and scenario combinations to assess the contribution of interest beyond usefulness and emotion. In one cohort, functional MRI examined age associations with responses during scenario reading that corresponded to subsequently reported appraisals, including within regions identified by a separate mental-state judgment task. These measures address age differences in adolescents’ choices as providers of social information.

## Results

### Subjective appraisals are more closely coupled to sharing with increasing age in adolescence

We examined gossip decisions in two independent samples aged 10–18 years: a behavioural sample (Sample 1, n = 98) and an fMRI sample (Sample 2, n = 101 scanned), of whom 100 provided complete outside-scanner behavioural data. Participants read scenarios about another person, decided whether to share the information and, if they chose to share, selected a recipient category. They then rated their interest in the information, its usefulness and their emotional response, alongside additional appraisals that differed between samples (Fig. 1). Sample 1 characterised behavioural age differences; Sample 2 examined whether these patterns were also observed in an independent sample and investigated their neural correlates. We first asked whether age was associated with the proportion of trials on which participants shared information and with valuation-to-decision coupling.

**Fig. 1.**
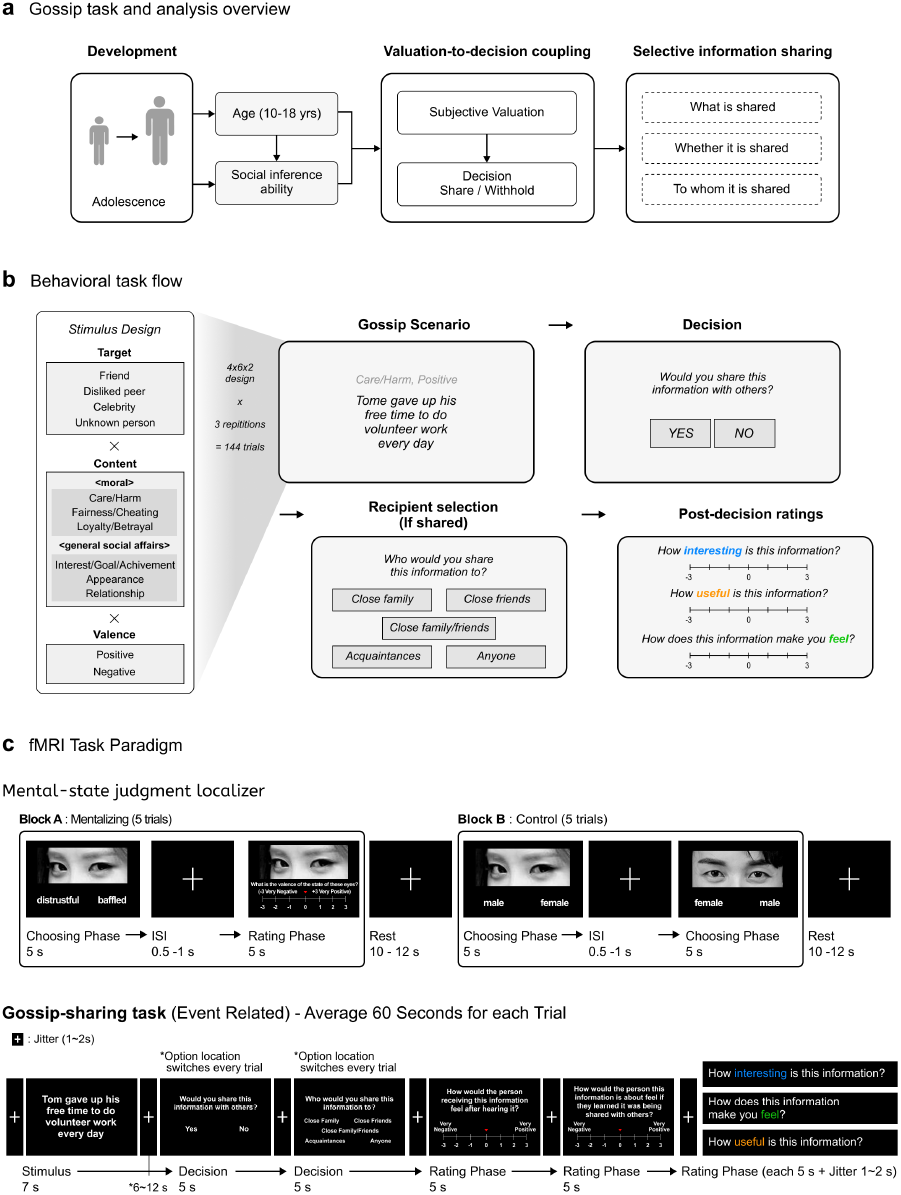
Conceptual framework and experimental design. **a,** Conceptual framework illustrating the proposed relationships among age, social inference ability, valuation-to-decision coupling, and the selection of information and recipients. Valuation-to-decision coupling describes how subjective appraisals relate to decisions to share or withhold social information. The diagram summarises the study hypotheses. **b,** Behavioural paradigm used in independent behavioural (Sample 1, *n* = 98) and fMRI (Sample 2, *n* = 101) cohorts aged 10–18 years. Participants read a gossip scenario, decided whether to share it and, if they chose to share, selected a recipient from five options. Ratings followed the sharing decision. The three appraisal dimensions common to both studies—interest, usefulness and emotion—are illustrated. Sample 1 additionally assessed information valence and its expected effect on the target; Sample 2 assessed the anticipated reactions of the recipient and target. The behavioural design crossed four target types, six content domains and two valences, with three repetitions per combination, yielding 144 trials. Sample 2 completed 84 out-of-scanner and 40 in-scanner trials. Social inference ability was assessed using the Korean Reading the Mind in the Eyes Test (K-RMET)^22^. **c,** In-scanner task sequence. Participants first completed a mental-state judgment localizer comprising alternating mental-state and sex-judgment blocks (A–B–A–B; five trials per block). Only mental-state judgments were followed by a valence rating. The localizer was analysed using an event-related model, with judgment and rating events modelled separately. Participants then completed 40 gossip-sharing trials. Each scenario was presented for 7 s, followed by a jittered 6–12-s interval before the sharing decision. The share/withhold decision, recipient selection following a decision to share, and each subsequent rating had a maximum response window of 5 s. The five ratings assessed recipient perspective, target perspective, interest, emotion and usefulness, in that order. Brief jittered fixations separated the remaining trial stages. Behavioural responses were self-paced. In the fMRI task, the 5-s labels denote maximum response windows; scenario presentation was fixed at 7 s.

We examined sharing frequency in Sample 1 (n = 98) and in the subset of Sample 2 with combined outside- and inside-scanner behavioural data (n = 92). Sharing frequency, defined as the proportion of valid trials on which a participant chose to share, was not detectably associated with age in either sample (Sample 1: r = 0.043, P = 0.671; Sample 2: r = −0.166, P = 0.114; Fig. 2a). The pooled correlation across samples was r = −0.058 (95% confidence interval [CI], −0.200 to 0.086; P = 0.427). Thus, neither sample provided clear evidence of an association between age and sharing frequency.

**Fig. 2.**
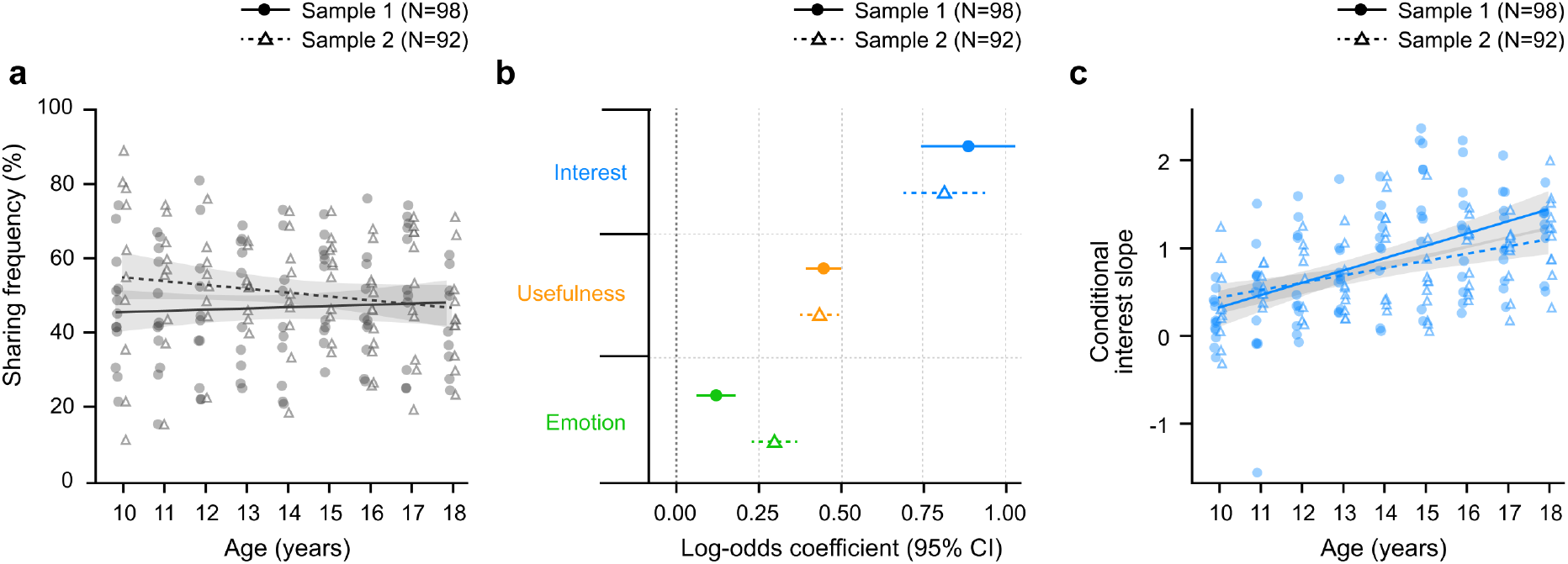
Subjective appraisals are more closely coupled to sharing with increasing age in adolescence. **a,** Percentage of valid trials shared against age in Sample 1 (n = 98) and the combined-task Sample 2 behavioural subset (n = 92), combining valid out-of-scanner and in-scanner trials for Sample 2. Each point represents one participant and lines indicate fitted linear trends. Sharing frequency was not detectably associated with age in either sample (Sample 1: r = 0.043, P = 0.671; Sample 2: r = −0.166, P = 0.114). The pooled association was r = −0.058, 95% CI [−0.200, 0.086], P = 0.427. **b,** Fixed effects of within-participant standardised interest, usefulness and emotion ratings on sharing, evaluated at the reference age in each sample (standardised age = 0, corresponding to the mean age used for age standardisation), from a joint mixed-effects logistic model including age-by-appraisal interactions, content and valence, participant and scenario random intercepts and participant random slopes for interest (Sample 1, n = 98; combined-task Sample 2 behavioural subset, n = 92). Symbols show log-odds coefficients and horizontal bars indicate 95% Wald confidence intervals. All three ratings were positively associated with sharing in both samples (all P < 0.001). **c,** Age-related difference in the association between interest and sharing in the same n = 98 and n = 92 samples. Points show conditional interest slopes from the mixed-effects model, combining the fixed interest effect, age-by-interest term and participant-specific random slope; these differ from the Bayesian weights in Fig. 3. Lines and shaded bands show descriptive linear fits and approximate 95% confidence intervals for the fitted mean; inference on the age-by-interest association was obtained from the mixed-effects model. The age × interest interaction was positive in Sample 1 (β = 0.377, 95% CI [0.235, 0.519]) and Sample 2 (β = 0.221, 95% CI [0.098, 0.343]) and pooled to β = 0.287, 95% CI [0.195, 0.380], P < 0.001. Circles and solid lines denote Sample 1; triangles and dashed lines denote Sample 2. Shaded bands in a and c and horizontal bars in b show 95% confidence intervals.

We next examined whether valuation-to-decision coupling, defined as the strength of the association between appraisal ratings and sharing choices, differed with age. We fitted logistic mixed-effects models to trial-wise sharing choices, including interest, usefulness and emotion ratings, their interactions with age, and content and valence covariates. The models accounted for differences in baseline sharing tendencies across participants and scenarios, and for individual differences in the association between interest and sharing decisions. At the mean age used to standardise age in each sample, higher interest, usefulness and emotion ratings were each associated with a greater probability of sharing (Sample 1, n = 98; Sample 2, n = 92; all P < 0.001; Fig. 2b; see Cross-cohort pooling in Methods). The age-by-interest interaction was positive in both samples (Sample 1: β = 0.377; Sample 2: β = 0.221; pooled β = 0.287, 95% CI, 0.195 to 0.380; P < 0.001; Fig. 2c). Thus, the association between interest and sharing decisions strengthened with age in both samples. The age-by-interest interaction did not differ detectably between samples (likelihood-ratio *χ*^2^(1) = 2.43, P = 0.119). The age-by-interest interaction remained positive in both samples in models adjusting for individual differences in rating-scale use and in separate models adjusting for sex (Supplementary Table 4).

The age-by-usefulness interaction was also positive in both samples (pooled β = 0.108, 95% CI, 0.068 to 0.149). The age-by-emotion interaction differed between samples (heterogeneity Q test, P = 0.043; Extended Data Fig. 1 and Supplementary Table 2). Together, these results indicated that the associations of interest and usefulness with sharing decisions strengthened with age in both samples, whereas the age-related difference in the association between emotion and sharing decisions varied between samples.

### Age is associated with stronger valuation-to-decision coupling and greater relative weight for interest

We next estimated participant-specific appraisal weights to characterise individual differences and examine their relationships with social-inference performance. We fitted the same hierarchical Bayesian model to both cohorts, including interest, usefulness and emotion simultaneously (Sample 1, n = 98; Sample 2, n = 100; Fig. 3a). Ratings were standardised within participant. The weights were log-odds coefficients relating each appraisal to sharing while accounting for the other two appraisals. The interest coefficient indexes valuation-to-decision coupling for interest, quantifying how strongly interest ratings are associated with sharing decisions. Both this model and the preceding mixed-effects model were fitted to trial-wise choices. We first estimated each participant’s interest, usefulness and emotion weights without including age in the model. We then examined whether these weights were associated with age. For Sample 2, these weights used out-of-scanner responses; the preceding mixed-effects model combined in-scanner and out-of-scanner responses.

**Fig. 3.**
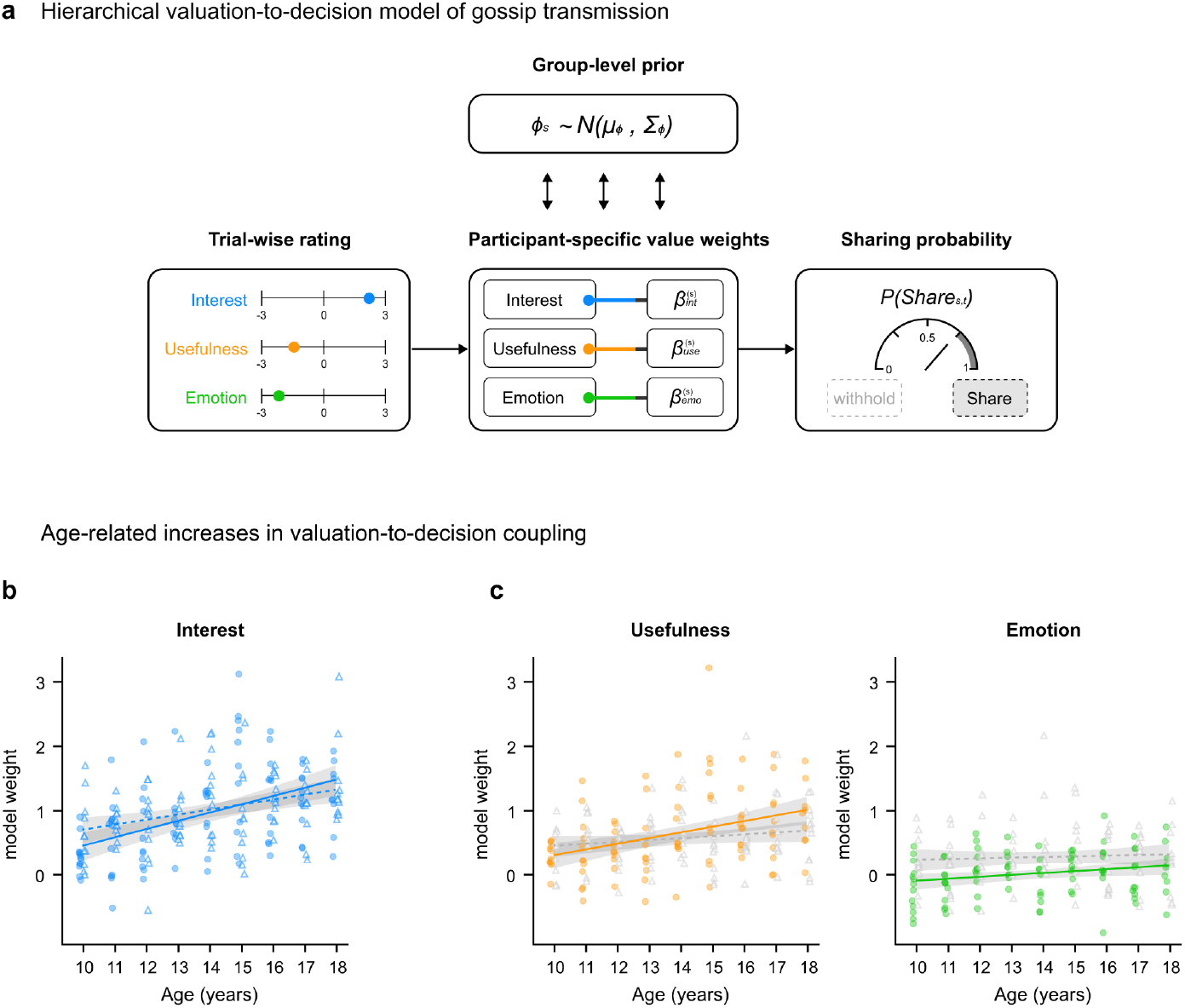
Participant-specific appraisal weights and their associations with age. **a,** Hierarchical Bayesian valuation model relating within-participant standardised post-decision interest, usefulness and emotion ratings simultaneously to sharing choices, with a participant-specific intercept and appraisal weights estimated under group-level priors. **b,** Interest weights were positively associated with age in Sample 1 (r = 0.475, 95% CI [0.306, 0.616], P < 0.001, n = 98) and Sample 2 (r = 0.351, 95% CI [0.167, 0.513], P < 0.001, n = 100). **c,** Usefulness weights were positively associated with age in Sample 1 (r = 0.376, P < 0.001) and showed a smaller same-direction association in Sample 2 (r = 0.169, P = 0.094). Emotion weights were positively associated with age in Sample 1 (r = 0.215, P = 0.033) but not Sample 2 (r = 0.055, P = 0.584). Appraisal weights are participant-specific posterior mean log-odds coefficients for within-participant standardised ratings, before decomposition into overall magnitude and relative components. Each weight indexes valuation-to-decision coupling for the corresponding appraisal. Each point represents one participant (Sample 1, n = 98, filled circles; Sample 2, n = 100, open triangles). Lines show ordinary least-squares fits and shaded bands indicate pointwise 95% confidence intervals for the fitted mean. Solid and dotted lines denote Samples 1 and 2, respectively. Coloured trends indicate uncorrected two-sided Pearson P < 0.05; other trends are shown in grey.

Interest weights were positively associated with age in both cohorts (Sample 1: r = 0.475, 95% CI [0.306, 0.616], P < 0.001; Sample 2: r = 0.351, 95% CI [0.167, 0.513], P < 0.001; Fig. 3b). The corresponding usefulness and emotion associations were less consistent across cohorts (Fig. 3c and Supplementary Table 4). These participant-level patterns were consistent with the positive age-by-interest interactions in the mixed-effects models and provided the individual-difference measures for the subsequent analyses.

Model validation assessed the recovery of participant-specific appraisal weights and the reproduction of sharing patterns across design cells. Simulated-data recovery reproduced the participant-specific ordering of appraisal weights, with median recovery correlations for interest of 0.916 and 0.828 in Samples 1 and 2. Fitted sharing probabilities also reproduced the observed pattern across the 48 target-by-content-by-valence cells (r = 0.953 and 0.939; Extended Data Fig. 2). Model comparisons supported the contribution of interest beyond usefulness and emotion (Supplementary Table 3).

Interest also improved out-of-sample prediction of sharing. We compared the three-appraisal model with a usefulness- and-emotion model using ten-fold cross-validation in Sample 1 (n = 98) and the combined-task Sample 2 behavioural subset (n = 92). One scheme held out entire participants; the other held out target-by-content-by-valence combinations. Both used post-decision appraisal ratings to predict sharing decisions that had not been used to fit the relevant model. Adding interest reduced held-out log loss, indicating lower prediction error, in all four sample-by-validation conditions. Mean participant-specific improvements were 0.023 and 0.032 for participant holdouts and 0.050 and 0.044 for design-cell holdouts in Samples 1 and 2, respectively (all P ≤ 0.0067; Extended Data Fig. 3). Interest therefore provided predictive information beyond usefulness and emotion across both participants and scenario combinations.

Finally, we asked whether larger interest weights reflected greater overall appraisal-weight magnitude, greater relative interest weight, or both. Overall magnitude was the Euclidean norm of the three-appraisal coefficient vector; relative interest weight was the interest coefficient divided by this magnitude. Overall magnitude captures the combined strength of the three appraisal coefficients, whereas relative interest weight captures the interest coefficient in relation to that overall magnitude. Overall magnitude was positively associated with age in both samples (pooled r = 0.444, 95% CI [0.324, 0.550], P < 0.001). Relative interest weight showed a positive pooled association with age (r = 0.246, 95% CI [0.110, 0.374], P < 0.001; Extended Data Fig. 4).

### Social inference covaries with valuation-to-decision coupling for interest across cohorts

We next examined how valuation-to-decision coupling for interest was related to social-inference ability, assessed with the Korean Reading the Mind in the Eyes Test (K-RMET), a test of mental-state recognition from the eye region. These analyses included 98 participants with complete K-RMET data in each cohort and used interest weights from the three-appraisal model. K-RMET performance was positively associated with age in both samples (r = 0.385 and 0.474, both P < 0.001; Fig. 4a). Higher K-RMET scores were also associated with larger interest weights (Sample 1: r = 0.402, P < 0.001; Sample 2: r = 0.311, P = 0.0018; Fig. 4b). Among the common appraisal weights, interest was the only dimension whose association with K-RMET reached significance in both cohorts (Supplementary Table 5).

**Fig. 4.**
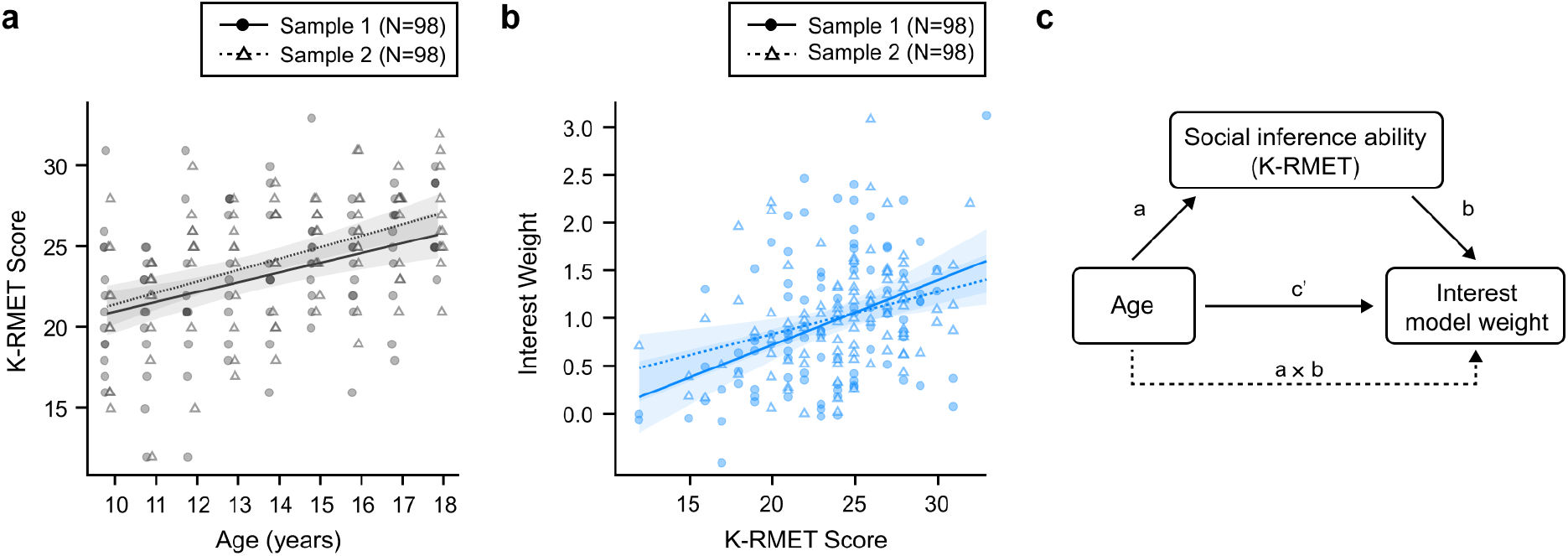
Social inference covaries with valuation-to-decision coupling for interest across cohorts. **a,** K-RMET performance was positively associated with age in Sample 1 (r = 0.385, P < 0.001) and Sample 2 (r = 0.474, P < 0.001). **b,** Higher K-RMET scores were associated with larger interest weights from the common three-appraisal model in Sample 1 (r = 0.402, P < 0.001) and Sample 2 (r = 0.311, P = 0.0018). Panels a and b include 98 participants per sample. Points represent participants; circles and solid lines denote Sample 1, and open triangles and dotted lines denote Sample 2. Lines show ordinary least-squares fits and shaded bands show pointwise 95% confidence intervals for the fitted mean. Age positions are jittered for visualisation only. **c,** Interest-related paths from the primary structural equation model, fitted separately in each sample (n = 98 each): a, age to K-RMET; b, K-RMET to interest weight conditional on age; *c*^!^, age to interest weight conditional on K-RMET; and a × b, the indirect association through K-RMET. Variables were standardised within each sample; additional model paths are described in Methods. Indirect point estimates were positive in both samples, but the 95% bootstrap confidence interval excluded zero only in Sample 1 (Sample 1: β = 0.094, 95% bootstrap CI [0.013, 0.192], P = 0.046; Sample 2: β = 0.077, 95% bootstrap CI [−0.031, 0.173], P = 0.122). Age retained a direct association with the interest weight in both samples (β = 0.382 and 0.294, respectively; both P < 0.004). Structural-model confidence intervals are percentile intervals from 1,000 case-bootstrap resamples; two-sided Wald P values use bootstrap standard errors. Paths represent cross-sectional associations.

Structural equation models estimated the associations among age, K-RMET and the interest weight jointly, distinguishing the direct age association from the indirect association through K-RMET (Fig. 4c). Additional appraisal-weight paths are specified in Methods.

Indirect point estimates were positive in both cohorts, but the 95% bootstrap confidence interval excluded zero only in Sample 1 (Sample 1: β = 0.094, 95% bootstrap CI [0.013, 0.192], P = 0.046; Sample 2: β = 0.077, 95% bootstrap CI [−0.031, 0.173], P = 0.122). Age retained a positive direct association with the interest weight in each cohort (β = 0.382 and 0.294, both P < 0.004; Supplementary Table 7). Thus, K-RMET performance was associated with the interest weight in both samples, and the age association remained after accounting for K-RMET. Comparisons with other cognitive and attachment measures and alternative structural specifications are reported in Supplementary Tables 6 and 7.

### Age is associated with what adolescents share and whom they choose to tell

We next examined age associations with whom adolescents chose to tell and which types of social information they chose to share (Sample 1, n = 98; Sample 2, n = 100). For each participant, we calculated the proportion of sharing decisions directed to each recipient category, conditional on choosing to share. The proportion of sharing decisions directed to close friends increased with age in both samples (Sample 1: r = 0.281, P = 0.005; Sample 2: r = 0.429, P < 0.001; pooled r = 0.358, 95% CI [0.229, 0.475], P < 0.001; Fig. 5a).

**Fig. 5.**
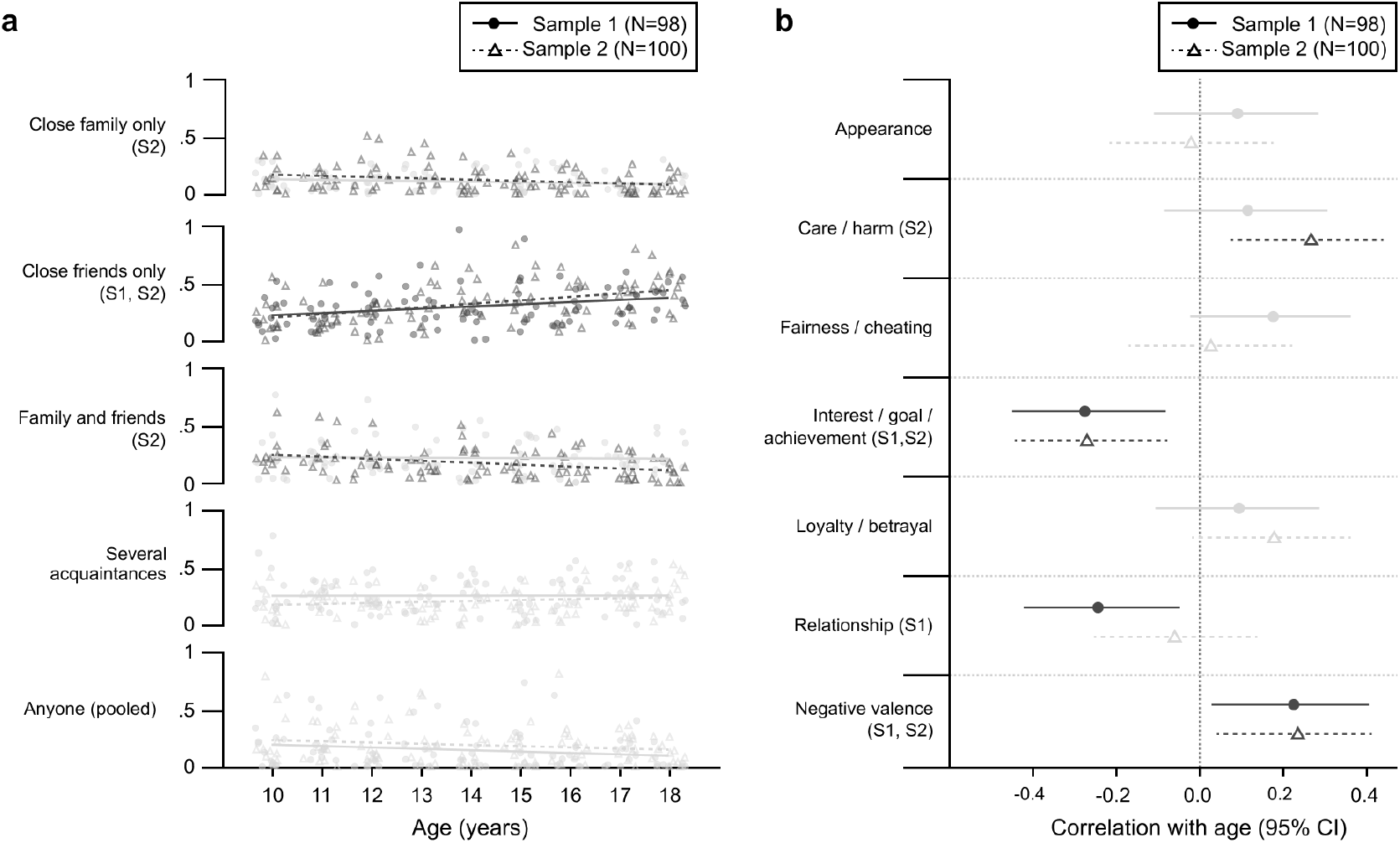
Age differences in whom and what is shared. **a,** Recipient selection among trials on which information was shared (Sample 1, n = 98; Sample 2, n = 100). Each proportion is the number of trials assigned to a recipient category divided by all trials on which that participant chose to share. Points represent participants and lines show ordinary least-squares fits; age positions are jittered for visualisation only. The proportion directed to close friends was positively associated with age in Sample 1 (r = 0.281, P = 0.005) and Sample 2 (r = 0.429, P < 0.001), with a pooled association of r = 0.358, 95% CI [0.229, 0.475], P < 0.001. Smaller pooled decreases were observed for close family (r = −0.208, P = 0.0034) and anyone including strangers (r = −0.172, P = 0.0158). **b,** Pearson correlations between age and participant-specific content-model weights in the same samples. The six content categories and negative valence are shown; coefficients were estimated jointly with four target indicators and both valence indicators using 12 within-participant mean-centred predictors. Symbols show correlations and horizontal bars indicate Fisher-z 95% confidence intervals. The weight for interest-, goal- and achievement-related content decreased in both samples (r = −0.273 and −0.268; pooled r = −0.270, FDR-adjusted q = 0.0015), whereas the weight for negatively valenced information increased (r = 0.228 and 0.238; pooled r = 0.233, q = 0.0041). Across panels, circles and solid lines denote Sample 1, and open triangles and dotted lines denote Sample 2. Dark marks and sample annotations identify within-sample associations with unadjusted two-sided P < 0.05; pale marks denote P ≥ 0.05. The pooled annotation refers to a pooled analysis rather than an additional plotted series. Pooled correlations used inverse-variance fixed effects on Fisher-z correlations, with weights n − 3; q values use Benjamini–Hochberg correction across all 12 pooled content-model tests. Of these two associations, only the interest-, goal- and achievement-related content association in Sample 2 survived the corresponding within-sample correction across 12 tests. Cohort-specific results are reported in Supplementary Table 9.

In pooled analyses, age was negatively associated with the proportions of sharing decisions directed to close family (r = −0.208, P = 0.0034) and to anyone, including strangers (r = −0.172, P = 0.0158). Recipient entropy, indexing how evenly sharing decisions were distributed across recipient categories, also showed a negative pooled association with age. However, an ordinal recipient-category score showed no clear age association (pooled r = −0.130, P = 0.070; Extended Data Fig. 5 and Supplementary Table 10). Together, these results indicate that, with increasing age, adolescents directed a greater proportion of their sharing decisions to close friends, a pattern observed in both samples.

Content-model coefficients revealed additional age-related differences in what was shared (Fig. 5b). These models estimated associations between content domains and sharing while including target and valence indicators. The coefficient for interest-, goal- and achievement-related content was negatively associated with age in both cohorts (r = −0.273 and −0.268; pooled r = −0.270, false-discovery-rate (FDR)-adjusted q = 0.0015). This category described the subject matter of a scenario and was distinct from participants’ ratings of how interesting they found the information. Conversely, the coefficient for negatively valenced information was positively associated with age in both cohorts (r = 0.228 and 0.238; pooled r = 0.233, q = 0.0041). These associations had the same direction and unadjusted P < 0.05 in each cohort; after correction across 12 tests within each cohort, only the interest-, goal- and achievement-related content association in Sample 2 remained significant. Both pooled associations survived correction across all 12 pooled content-model tests. An increase in care- and harm-related content weighting was concentrated in Sample 2 (Supplementary Table 9).

We also asked how appraisal weights and recipient selection were related across participants. A larger interest weight was associated with a lower proportion of sharing choices directed to anyone, including strangers, after adjustment for age in both cohorts (partial r = −0.228 and −0.261; pooled partial r = −0.245, 95% CI [−0.373, −0.107], P < 0.001; Supplementary Table 11). Thus, the association between the interest weight and this recipient preference was also evident after accounting for their respective relationships with age.

Having examined appraisal weights, recipient selection and content preferences separately, we asked whether these measures could be summarised by a single underlying dimension. A one-factor model fit poorly in both samples (CFI = 0.000 and 0.039). The first principal component showed similar loading patterns across samples (absolute Tucker’s congruence coefficient |*φ*| = 0.981), explaining 30.0% and 30.6% of the variance, respectively (Extended Data Fig. 6). A similar principal component emerged across samples, but the one-factor model fit poorly in both.

### Age associations with interest-related BOLD modulation during scenario reading

In the fMRI sample, we examined whether the association between BOLD responses during scenario reading and subsequently reported interest varied with age. In an initial model that included interest as the sole appraisal-related parametric modulator, interest-related BOLD modulation during scenario reading was positively associated with age in the left middle frontal gyrus (n = 97; peak MNI [−54, 18, 39]; k = 196; T = 4.46; cluster-level PFWE = 0.000577; Fig. 6a).

**Fig. 6.**
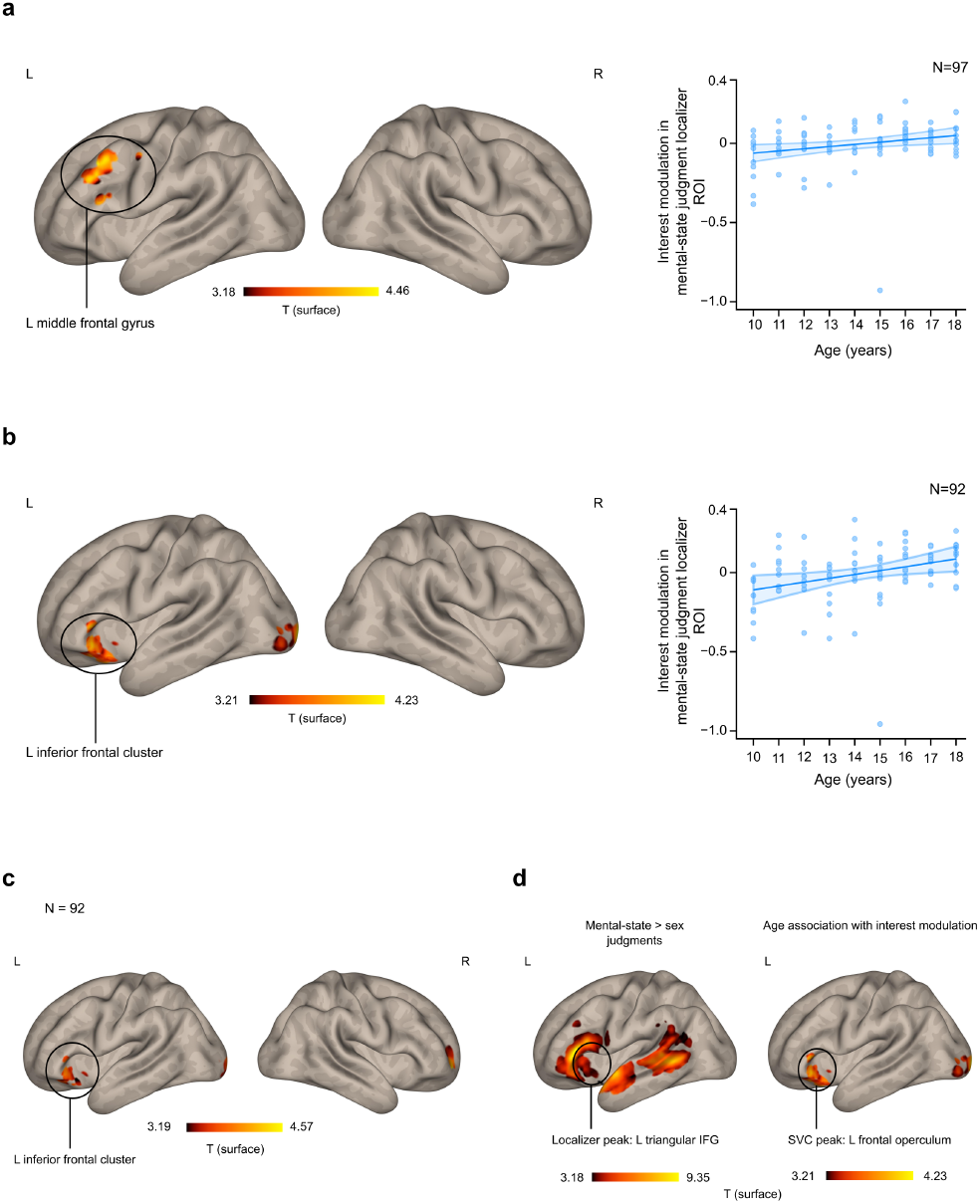
Age associations with interest-related BOLD modulation during scenario reading. **a,** In the initial analysis, scenario-interest modulation was positively associated with age in left middle frontal gyrus (n = 97; MNI [−54, 18, 39]; k = 196; T = 4.46; cluster-level PFWE = 0.000577). The plot shows mean raw scenario-interest modulation in an 8-mm sphere centred on the separate-task group-mean localizer peak (MNI [−57, 27, 3]), selected without an age regressor (age coefficient = 0.0139 ROI-estimate units per year, 95% CI [0.00288, 0.02492], nominal two-sided P = 0.0140). **b,** In the 92-participant joint model, interest, usefulness and emotion were entered simultaneously without serial orthogonalisation. The scenario-interest age association remained in a left inferior frontal cluster after adjustment for K-RMET, mean framewise displacement and sex (MNI [−39, 30, −9]; k = 116; T = 4.73; cluster-level PFWE = 0.000316). The plot shows partial residuals plus the fitted intercept and centred-age contribution in the same sphere, with covariates held at sample means (age coefficient = 0.0244 ROI-estimate units per year, 95% CI [0.00511, 0.04373], nominal two-sided P = 0.0138). **c,** A separately estimated model (n = 92) that additionally included the eventual share-or-withhold choice retained an age association at the same peak after adjustment for K-RMET and mean framewise displacement (MNI [−39, 30, −9]; k = 84; T = 4.80; cluster-level PFWE = 0.00199). **d,** Regions were defined by the group-mean contrast of mental-state versus sex judgments in a separate task (n = 99), without an age regressor. Small-volume correction used the full significant group-mean localizer network. The indicated peak from the final joint scenario-interest age map (n = 92) was significant within this mask (peak MNI [−45, 24, −3]; T = 4.48; peak-level PFWE-SVC = 0.0157). The scatterplots in a and b use the same separate-task localizer sphere, rather than the displayed whole-brain clusters. Each point represents one participant; lines show fitted conditional means and shaded bands show pointwise 95% confidence intervals derived from the full-model covariance. Surface maps were displayed at voxel P < 0.001 uncorrected; reported clusters passed whole-brain cluster-level FWE P < 0.05. Colour scales show surface-display T statistics. The ROI P values in a and b are nominal descriptive follow-ups and are not imaging FWE-corrected tests.

To examine whether the age association with interest-related BOLD modulation remained after accounting for usefulness and emotion, we entered all three appraisals simultaneously, without serial orthogonalisation. This analysis included 92 participants with valid interest contrasts for both scenario reading and sharing decisions, with the three appraisal modulators included for each phase. After adjustment for K-RMET, mean framewise displacement and sex at the group level, interest-related BOLD modulation during scenario reading was positively associated with age in a left orbital inferior frontal cluster extending into the frontal operculum and posterior orbital gyrus (peak MNI [−39, 30, −9]; k = 116; T = 4.73; cluster-level PFWE = 0.000316; Fig. 6b).

In an additional ROI analysis of the initial interest-only model, an association was also present within a fixed left lateral orbitofrontal sphere (observed peak MNI [−33, 27, −15]; T = 4.31; peak-level PFWE-SVC = 0.00062; Extended Data Fig. 7b).

We then asked whether the age association remained after accounting for the subsequent sharing choice, adding share-or-withhold choice to a separate model that also included interest, usefulness and emotion. After adjustment for K-RMET and mean framewise displacement at the group level, interest-related BOLD modulation during scenario reading was positively associated with age, with the same peak coordinate as in the three-appraisal fMRI model (n = 92; peak MNI [−39, 30, −9]; k = 84; T = 4.80; cluster-level PFWE = 0.00199; Fig. 6c).

Given the association between social-inference performance and valuation-to-decision coupling for interest in both cohorts, we examined whether age-related differences in interest-related BOLD modulation were also evident within regions defined by the mental-state judgment localizer. We defined these regions using the group-mean contrast of mental-state versus sex judgments in 99 participants, without an age regressor, and used the resulting mask for small-volume correction of the age association in the three-appraisal fMRI model.

Within this mask, interest-related BOLD modulation during scenario reading was positively associated with age (n = 92; peak MNI [−45, 24, −3]; T = 4.48; peak-level PFWE-SVC = 0.0157; Fig. 6d and Extended Data Fig. 8). The age association was also significant after small-volume correction within an 8-mm sphere centred on the group-mean localizer peak (centre MNI [−57, 27, 3]; observed peak MNI [−54, 21, 0]; T = 3.21; peak-level PFWE-SVC = 0.0404). The scatterplots in Fig. 6a,b show participant-level estimates from this localizer sphere; the adjacent brain maps show the corresponding whole-brain results. These findings place the interest-related age association within cortex also engaged during mental-state judgments.

In the initial interest-only model, the age association with interest-related BOLD modulation differed between scenario reading and sharing decisions (Extended Data Fig. 7c). This between-phase difference was not statistically significant when interest, usefulness and emotion were modelled simultaneously. An additional permutation analysis correcting across the three interest-related contrasts and both directions yielded no significant clusters (Supplementary Table 12).

## Discussion

Across two independent cohorts, sharing decisions were more closely related to reported interest with increasing age, and close friends accounted for a greater proportion of recipient choices. Sharing frequency showed no consistent age association across cohorts. Computational analyses distinguished stronger valuation-to-decision coupling overall from a greater relative contribution of interest in the pooled analysis, while social-inference performance was related to the interest weight in both cohorts. In the fMRI sample, BOLD responses during scenario reading were more strongly related to subsequently reported interest with increasing age. Together, these findings identify age-related differences in adolescent gossip in the correspondence between appraisals and sharing choices and in the selection of recipients.

The computational findings clarify what it means for interest to matter more for sharing at older ages. Previous research has shown that information can acquire value through both instrumental and non-instrumental motives, and that decisions to inform others reflect the anticipated consequences for recipients’ actions, feelings and knowledge^13,16^. In our task, interest contributed to sharing alongside usefulness and emotion, and the positive age-by-interest association recurred across both cohorts. The larger interest weights reflected two distinguishable patterns: a greater overall magnitude of appraisal coefficients in both samples and a greater relative contribution of interest in the pooled analysis. Interest also improved held-out prediction beyond usefulness and emotion.

We also examined whom adolescents chose to tell once they decided to share. Research on adolescent information seeking and social learning highlights uncertainty about others’ trustworthiness and preferences for particular information sources^23,24,25^. These findings complement research on whom adolescents choose as sources of information by examining whom they select as recipients of gossip. Close friends accounted for a greater proportion of sharing choices with increasing age in both cohorts. Moreover, a larger interest weight was associated with a lower proportion of sharing directed to anyone, including strangers, after accounting for age in both cohorts. Appraisal weights and recipient selection therefore showed a specific, reproducible relationship, connecting subjective appraisals of information with the audiences selected to receive it. These conditional proportions do not establish an increase in the absolute amount shared with close friends or a general narrowing of the recipient range.

Because gossip is directed to other people, an important question is whether valuation-to-decision coupling is related to the ability to infer others’ mental states. Computational accounts of mentalizing describe how people infer others’ beliefs and desires, and experimental evidence shows that evaluative contexts can facilitate mental-state inference^18,26^. Here, higher K-RMET performance was associated with larger interest weights in both cohorts, linking performance on this task with how reported interest relates to sharing. Although K-RMET performance was associated with interest weights in both samples, the age-adjusted associations were less consistent. The indirect association through K-RMET was supported only in Sample 1, and a direct age association remained in both samples. One possibility is that appraising information for sharing draws on both its interest to oneself and an understanding of its potential relevance to others. This account connects subjective appraisal to social understanding while allowing other experiences and motivations to contribute to age differences. K-RMET measures mental-state recognition from the eye region and does not capture all aspects of mentalizing in everyday social interaction.

Previous work has examined neural representations of social value during peer evaluation in adolescents^27^. Here, we examined interest-related responses while adolescents read social information they could subsequently share. A positive age association with interest-related BOLD modulation was observed during scenario reading, before participants reported their sharing decisions, linking age to neural responses to social information at a stage preceding the overt choice. The association persisted when interest, usefulness and emotion were modelled simultaneously without serial orthogonalisation, and a separate model retained it after explicitly representing eventual choice. Complementary models also accounted for social-inference performance, head motion and sex.

The frontal location and mental-state judgment localizer provide a basis for interpreting this pattern. Anterior ventrolateral prefrontal cortex has been implicated in retrieving social knowledge for judgments about others, and lesion evidence links left inferior frontal cortex to the ability to infer mental states from the eyes^28,29^. These findings provide a task-based context for interpreting the three-appraisal fMRI association in the left inferior frontal cluster and the SVC finding in left frontal operculum. Crucially, our functional reference came from a separate mental-state judgment task, with the localizer defined without an age regressor. The neural findings complement this behavioural association by locating age-related differences in interest-related BOLD modulation within regions also engaged during mental-state judgments. These observations motivate the hypothesis that social inference contributes to appraising information for sharing, without establishing a mediating mechanism.

The neural findings also require qualification. No cluster survived the separate synchronized permutation analysis across the three joint-model interest maps and both directions, and the direct phase contrast was not significant in the joint appraisal model. The within-map findings and the broader permutation analysis used different covariate specifications and correction families. The data therefore do not establish that the age association is specific to scenario reading or to a particular mental process.

Interest in a piece of information should also be distinguished from its subject matter. Previous work demonstrates that the target, subject matter and valence of social information jointly shape gossip decisions^21^. Here, the age-related increase in the weighting of subjective interest coexisted with decreased weighting of scenarios about interests, goals and achievements, and increased weighting of negatively valenced information. These content associations were less robust under multiplicity correction. What information is about and how interesting an individual finds it are distinct features of a sharing decision. The poor fit of the one-factor model suggests that these measures cannot be adequately summarised by a single latent dimension. It does not establish that the domains are independent or confirm a specific alternative factor structure.

These findings motivate longitudinal and experimental extensions. Replication of the interest– sharing association and its age dependence across two independent samples strengthens the evidence for this pattern during adolescence. These findings concern differences across ages; longitudinal studies could establish how this pattern unfolds within individuals. Because appraisal ratings were collected after sharing choices, the direction of the relationship between appraisals and sharing cannot be established. Varying information value and rating timing would help distinguish these possibilities. The scenarios measured intended sharing rather than actual information transmission or its social consequences. Reciprocal exchanges across cultural settings could establish how the observed patterns operate when sharing has consequences for relationships and reputations. Such extensions connect the present account of sharing decisions to evidence that gossip supports vicarious learning, social connection and cooperation^3,4^.

Gossip provides a window onto adolescents’ active role in social information exchange. Across cohorts, age differences concerned how strongly sharing decisions were coupled to subjective appraisals and whom adolescents selected to receive information. The corresponding neural age association was observed during scenario reading, including within frontal regions defined by the mental-state judgment localizer. Together, our findings broaden our understanding of adolescent social development from how young people respond to social information to how they appraise what is worth sharing and choose whom to tell.

## Methods

### Participants

We studied two independent samples of adolescents aged 10–18 years. Sample 1 (the behavioural study) completed the gossip-sharing task as a computer-based behavioural session. Sample 2 (the fMRI study) performed a modified version of the same task, administered in two parts: a behavioural session outside the scanner and a shortened, event-related version during functional MRI. The task was adapted for Sample 2, with the appraisal-rating set revised and the trial count reduced for scanning.

Sample 1 comprised 98 adolescents (mean age = 13.82 years, s.d. = 2.62, range 10–18; 46 male, 52 female). Sample 2 comprised 101 scanned adolescents (mean age = 14.08 years, s.d. = 2.61; 51 male, 50 female). Two participants were excluded from neuroimaging analyses because at least 40.0% of their valid framewise-displacement samples exceeded 0.5 mm, leaving 99 participants. The primary scenario–decision phase comparison included 92 participants with valid scenario-interest and decision-interest contrasts under the analysis-specific exclusion criteria and complete age, K-RMET, sex and mean-framewise-displacement covariates. Analysis-specific sample sizes are reported throughout. Both samples were recruited through online participant-recruitment advertisements, parenting forums and social media to enrol adolescents across the 10–18-year age range. Sample 1 data were collected from November 2024 to May 2025, and Sample 2 data were collected from June 2025 to January 2026. Exclusion criteria were a reported neurological or psychiatric disorder, current psychotropic medication use or uncorrected sensory impairment that would interfere with the task. Sample 2 additionally met standard MRI safety criteria and was screened for claustrophobia and metal implants.

#### Sampling strategy and demographic variables

We targeted approximately 10 participants per one-year age bin from 10 to 18 years, with an approximately balanced distribution of sex assigned at birth within each bin. The 10–18-year range was selected to span the proposed preadolescent transition from overt tattling to increasingly peer-oriented and covert forms of gossip, theorised to occur around 10–11 years^9,11^, through late adolescence. Sex was measured as participant-reported sex assigned at birth (male or female); gender identity was not assessed. Handedness was assessed by self-report; all participants in both samples reported being right-handed.

Participants received 25,000 KRW for the behavioural session; those in Sample 2 received 70,000 KRW upon completion of the MRI session, contingent on completing the full protocol. The study was approved by the institutional review board (IRB) of the Korea Advanced Institute of Science and Technology (KH2024-229). Written informed consent was obtained from a parent or legal guardian, and written assent from each adolescent participant, before participation.

The study was not preregistered. Random allocation to experimental groups and investigator blinding were not applicable because there was no group assignment and outcomes comprised computer-recorded choices, ratings and MRI signals.

### The gossip-sharing task

#### Overview and operational definition

The task was implemented in PsychoPy v2024.2.4^30^. The task was adapted from the adult gossip-sharing paradigm developed by Lee et al.^21^, with modifications for adolescent participants and the present study aims. On each trial, participants read a short scenario describing an action attributed to a specific individual, decided whether to share the information, and if so, selected a recipient. They then rated the scenario on several appraisal dimensions. We operationally defined valuation-to-decision coupling as the within-participant trial-level association between appraisal ratings and share/withhold choices. Participants were instructed that the task concerned transmitting information about an absent third party, the standard operational definition of gossip^1^. The instructions and on-screen prompts used neutral wording (“share this information with others”) and avoided the evaluative Korean term for gossip (*dwitdamhwa*), so that its negative connotations would not bias sharing decisions. The same framing was used in both samples.

#### Trial structure

Each scenario was presented for 7 s; participants were instructed to treat it as a real event and consider whether to share it. Subsequent responses were self-paced in the behavioural sessions; MRI response windows are specified below. They then made a binary share/withhold decision in response to the prompt, “Would you share this information with others?” If they chose to share, they selected a recipient option from five categories: close family, close friends, family and friends, several acquaintances, or anyone including strangers. The “anyone” category explicitly included strangers and sharing in online or first-encounter contexts. Following the sharing decision, participants rated the scenario on a set of appraisal dimensions, each on a 7-point Likert scale from −3 to +3.

#### Appraisal ratings

Three appraisal dimensions were common to both samples and form the basis of cross-sample appraisal analyses: *interest* (“How interesting is this information?”; −3 = very boring, +3 = very interesting), *emotion* (“How does this information make you feel?”; −3 = very bad, +3 = very good), and *usefulness* (“Is this information useful – worth knowing for yourself or others?”; −3 to +3). Informed by the initial study, the remaining two ratings were refined for Sample 2. Sample 1 additionally rated the *valence* of the information (whether its content was positive or negative, independent of the target) and the likely *effect on the target* were information to become known. In Sample 2, this single target-directed judgment was separated into two perspective ratings: the anticipated emotion of the recipient upon learning the information (*recipient perspective*) and of the *target* upon learning that the information was being shared (*target perspective*). Throughout, we refer to the five Sample 2 appraisal dimensions as interest, emotion, usefulness, target perspective, and recipient perspective. Because only interest, emotion, and usefulness were measured identically in both samples, all cross-sample replication analyses are restricted to these three dimensions; sample-specific ratings are reported within their respective samples.

#### Stimuli and personalisation

Scenarios spanned three moral domains (care/harm, fairness/cheating and loyalty/betrayal) and three general social domains (relationship, interest/goal/achievement and appearance). Each contained positive- and negative-valence versions, forming a fully crossed content × valence design (the shortened in-scanner task used five of these domains, omitting relationship). Each scenario concerned one of four target types defined relative to the participant: a liked close friend, a disliked peer or acquaintance, a famous celebrity the participant neither liked nor disliked, and an unfamiliar individual unknown to the participant. At the start of the session, participants supplied the real names of people in the first three categories (a liked close friend, a disliked peer or acquaintance, and a famous-but-neutral celebrity); Sample 1 provided three names per category and Sample 2 one. On each trial, the relevant name was inserted into the scenario so that it referred to a specific person in the participant’s social environment. The unfamiliar-target condition used a real name. Participants were told that the named person was unfamiliar to them, and the experimenter confirmed that they did not know anyone by that name.

#### Implementation and differences between samples

In Sample 1, responses were self-paced and made using a keyboard; participants completed 144 trials (4 targets × 6 content domains × 2 valences × 3 repetitions). Sample 2 completed 124 trials in total: 40 in the event-related version administered during scanning (*see fMRI task and acquisition*) and 84 in an out-of-scanner version with self-paced responses. The in-scanner version was reimplemented with fixed response windows. Within Sample 1, the 48 target × content × valence combinations were randomised separately in each of three repetitions, and scenarios were sampled without replacement within content × valence pools. In Sample 2, trial order was randomised separately for each participant. In both the out-of-scanner and in-scanner Sample 2 tasks, the left–right positions of the share/withhold and recipient options were randomised on every trial. Participants made responses predominantly with the right hand during the self-paced computer sessions and used both hands during the MRI task.

### Psychometric measures

#### Social inference

Social inference ability was indexed with the Korean version of the Reading the Mind in the Eyes Test (K-RMET)^22,31^. On each of 36 items, participants viewed a photograph of the eye region of a face and chose which of four mental-state terms best described what the person was thinking or feeling. The total number of correct responses served as the index of social inference ability.

#### Additional measures

The additional questionnaires are described in the Supplementary Methods; their descriptive statistics and internal consistency are reported in Supplementary Table 13.

### Behavioural analysis

#### Cross-sample testing

The samples shared interest, emotion, and usefulness ratings, the sharing decision, and recipient choice, but differed in their additional ratings. Cross-sample tests therefore used only the common measures. We first estimated each effect within each cohort, then combined comparable estimates by fixed-effect pooling. Trial-level age × rating interactions were additionally tested in a joint mixed model containing sample × age × rating terms; likelihood-ratio tests assessed whether age-related slopes differed between cohorts.

#### Sharing frequency and single-appraisal analyses

For each participant we computed the sharing proportion as the number of share responses divided by the number of valid share/withhold responses and related it to age using Pearson correlation. Figure 2a uses Sample 1 (n = 98) and the combined-task Sample 2 behavioural subset (n = 92), combining valid out-of-scanner and in-scanner responses in Sample 2. We then related each of the three common ratings – interest, emotion, and usefulness – to the binary share/withhold decision using single-predictor logistic regressions. Single-predictor results are reported in Supplementary Table 2. The joint mixed-effects model used for Fig. 2b,c is described below. Additional sample-specific ratings entered the hierarchical models.

#### Hierarchical Bayesian models of sharing

We examined three behavioural domains: subjective valuation of the information, recipient choice, and shared content. Each was modelled with hierarchical Bayesian logistic regression under a mixed-effects scheme using MATLAB and the Variational Bayesian Analysis (VBA) toolbox (commit 5899497f)^32^, which jointly estimates participant-level posteriors and group-level priors. The domains used related but distinct outcome and design matrices and were not assumed to reflect one latent construct. The model allowed the number of valid trials to vary across participants and retained all participants who met the data-inclusion criteria for each analysis.

The *valuation model* estimated the weight linking each appraisal rating to the probability of sharing. For participant *i* on trial *t*, the choice to share, *y_it_* ∈ {0,1}, was a Bernoulli outcome whose probability was a logistic function of a participant-specific intercept and a weighted sum of that trial’s appraisal ratings:

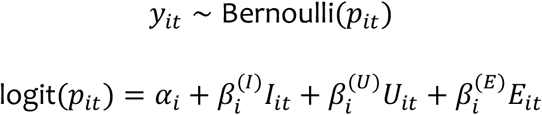

Where *I_it_*, *U_it_*, and *E_it_* denote trial-wise ratings of interest, usefulness, and emotion, respectively, and *α_i_* is a participant-specific intercept. Individual parameters were assumed to arise from a group-level distribution,

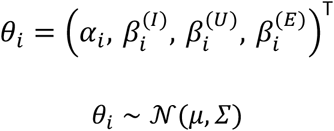

Participant-specific estimates were regularised by the population prior. The parameters 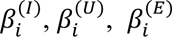 were interpreted as participant-specific valuation-to-decision weights; larger positive values indicated stronger associations between appraisal ratings and sharing. The three-appraisal model shown above was fitted separately in Sample 1 (n = 98, 14,113 valid trials) and the outside-scanner Sample 2 dataset (n = 100, 8,400 trials) and supplied the weights used in cross-cohort analyses. Cohort-specific full models additionally included the study-specific ratings available in each sample and were evaluated separately.

The *content model* used hierarchical logistic regression with 12 participant-centred indicator variables describing each scenario and no separate intercept:

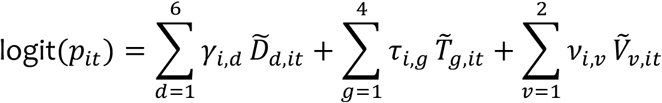

where 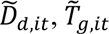 and 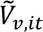 are indicator variables for the six content domains, four target types and two valences, respectively, each centred by subtracting its participant-specific mean. All 12 indicators entered the model; no reference category was omitted. The corresponding participant-specific weights were related to age across participants.

The *category-specific recipient-choice models* were fitted separately for each of the five recipient categories. The binary outcome was 1 when a participant chose to share with category *k* and 0 for all other trials, including withheld trials. For each category, the logistic model included a participant-specific intercept and a cohort-specific design matrix of appraisal ratings, selected absolute-rating terms, target indicators and selected appraisal-by-target interactions. The interactions involved target-effect ratings in Sample 1 and target- and recipient-perspective ratings in Sample 2, rather than all appraisal-by-target combinations. Associations between the category-specific intercepts and age are reported as supplementary recipient-model analyses in Supplementary Table 8; the main recipient-composition results use observed proportions conditional on sharing.

As in the valuation model, participant-level parameters in the content and recipient-choice models were drawn from group-level Gaussian priors, and models were specified separately by sample. Model-specific results are reported in Supplementary Tables 8 and 9. Cross-sample recipient-composition analyses used the observed proportion of shared trials assigned to each of the five categories rather than treating the categories as a single ordinal continuum.

#### Model validation

The three-appraisal model was evaluated by parameter recovery and comparisons of observed and fitted sharing rates. The close-friend and anyone recipient models were additionally refitted using reference-cell coding and evaluated by comparing observed and fitted choice rates and by a single simulation-and-refit assessment of coefficient recovery.

Recipient-choice findings were additionally evaluated for convergence across hierarchical, mixed-effects and distributional analyses. Free-energy comparisons of models with and without interest quantified its in-sample contribution. Split-half and participant-resampling reliability estimates are reported in Supplementary Table 14. The common three-appraisal model comparisons and validation analyses were run in MATLAB R2023b Update 11.

#### Trial-level mixed-effects models

To estimate age-by-appraisal associations directly while accounting for participant- and scenario-level variation, we used mixed-effects logistic regression (glmer in R, lme4 v1.1-37)^33^, predicting the trial-wise share/withhold choice. For valuation, age (z-scored) was entered in interaction with each of the three common ratings, with content and valence as fixed-effect factors. The models included participant and scenario random intercepts and participant random slopes for interest; the age × interest interaction provided a direct test of age differences in the interest–sharing association. The joint appraisal model was fitted in R v4.5.0 to 14,113 valid trials from Sample 1 (n = 98) and 11,347 valid out-of-scanner and in-scanner trials from Sample 2 (n = 92). Trials without a valid share/withhold response were excluded. To examine whether this effect was robust to age-related differences in rating-scale use, we refit the valuation model with three per-participant scale-use covariates (the standard deviation, range, and proportion equal to the participant’s rounded median rating, computed from ratings pooled across the three dimensions) and an interest × scale-use (s.d.) interaction. Parallel trial-level models tested the age × content interaction (reference: care/harm) and age-related differences in recipient choice (close-friends vs. other recipients; and an ordinal cumulative-link model of recipient breadth).

#### Cross-cohort pooling

The trial-level model was fitted separately by cohort with ratings z-scored within participant. Age was z-scored within each sample using the trial-level dataset before the final complete-case filter. Accordingly, the appraisal main-effect coefficients describe valuation-to-decision coupling at the sample-specific mean age used for standardisation, where standardised age equals zero. Comparable coefficients were combined by fixed-effect pooling. We then fit a joint model containing sample × age × rating interactions, with cohort-specific scenario identifiers, and used a likelihood-ratio test of sample × age × interest to assess slope heterogeneity. Separate extensions of the baseline model tested robustness to scale-use covariates, quadratic-age terms, and sex plus its interaction with interest. Within each cohort, these sensitivity models used the same valid-response trials, scenario identifiers and standardised age and appraisal variables as the primary model (Sample 1: 14,113 trials, n = 98; Sample 2: 11,347 trials, n = 92); scale-use covariates were calculated from those valid-response trials. These sensitivity models were fitted in R v4.5.2 with lme4 v1.1-37.

#### Social inference and structural-equation models

We related K-RMET performance to age and to participant-specific appraisal weights using Pearson correlations. To ensure that the interest weight was defined identically across cohorts, structural equation models (R v4.5.0, lavaan v0.6-19^34^) used interest weights from the model including interest, usefulness and emotion and were fitted separately in each cohort (n = 98 per cohort). The models estimated the indirect association between age and the interest weight through K-RMET while retaining a direct age-to-interest path. The primary specification also included paths from age to the usefulness and emotion weights, a path from K-RMET to the emotion weight, and residual covariance between the interest and usefulness weights. Models used integer age and maximum-likelihood estimation. Percentile-bootstrap 95% confidence intervals based on 1,000 resamples were the primary inferential criterion for the indirect association; corresponding two-sided z-based P values are also reported. Alternative path specifications, including a saturated model, assessed robustness (Supplementary Table 7). Indirect effects were interpreted as cross-sectional associations.

#### Out-of-sample prediction and weight decomposition

We compared an intercept-only model, a usefulness-plus-emotion model, the same model with interest added, and the sample-specific full model using 10-fold cross-validation (seed = 42) in Sample 1 (n = 98) and the combined-task Sample 2 behavioural subset (n = 92, 11,347 valid trials). Sample 2 cross-validation combined out-of-scanner and in-scanner trials, whereas the participant-specific weights from the three-appraisal model above used the out-of-scanner dataset. Participant-level folds held out entire participants and predicted them from the training group posterior. Scenario-level folds held out target × content × valence cells, applied training-set rating standardisation, and predicted held-out cells from each participant’s posterior. Log loss (negative log predictive probability) was primary; Brier score, held-out log predictive density and accuracy were secondary. To separate the overall magnitude of the appraisal weights from their relative weighting, each participant’s three-rating vector β was decomposed as β = γw, where γ = || β|| and w = β/γ.

#### Distribution of recipient choices and cross-domain structure

For each participant, we calculated the proportion of sharing choices directed to each of five recipient categories, Shannon entropy under three estimators, and an ordinal recipient-category score based on the ordered options (1 = close family only; 5 = anyone, including strangers). Comparable correlations with age were pooled across cohorts on the Fisher-z scale. To test whether valuation, recipient, and content measures formed one construct, we compared one-factor and three-factor confirmatory models, assessed the similarity of principal-component loading patterns across samples using Tucker’s congruence coefficient^35^, and tested cross-domain participant-level correlations. Partial correlations adjusted for age; the complete raw and age-adjusted matrix is reported in Supplementary Table 11. Minimum-sharing-trial sensitivity is reported in Supplementary Table 10. Because the three-factor model also showed imperfect absolute fit, these analyses assessed the specified one-factor model rather than validating a definitive alternative measurement model.

### fMRI task and acquisition

#### Task during scanning

In Sample 2, participants completed a shortened, fixed-timing version of the sharing task inside the scanner, structured as an event-related design. Each trial began with a jittered fixation cross (1–2 s), followed by the scenario presented for 7 s. After a jittered delay (6– 12 s) that allowed the scenario and decision events to be separated, participants made the share/withhold decision within a 5 s window; if they chose to share, they then selected a recipient (5 s window). They then provided the appraisal ratings, each within a 5 s window, separated by brief jittered fixations (1–2 s). Participants completed 40 scenario trials (4 targets × 5 content domains × 2 valences).

#### Mental-state judgment localizer

Before the sharing task, participants completed a mental-state judgment localizer adapted from the Reading the Mind in the Eyes paradigm^22,31,36^, alternating between a mental-state judgment condition and a sex-judgment control condition. On each trial, participants viewed a photograph of the eye region of a face for up to 5 s. In the mental-state judgment condition, they chose which of two words better described the person’s mental state; in the sex condition, they judged whether the eyes belonged to a man or a woman. Mental-state judgment trials were followed by a valence rating (5 s). The 20 trials were arranged in a fixed alternating sequence of mental-state judgment and sex-judgment sets (five trials per set; mental-state judgment–sex judgment–mental-state judgment–sex judgment), with trial order randomised within each set. Each set was preceded by a 2 s cue indicating the upcoming condition, with inter-trial fixations of 0.5–1.0 s and 10–12 s rest between sets. The fMRI analysis was event-related: the stimulus-judgment event and the subsequent valence-rating event (present only in the mental-state judgment condition) were modelled as separate regressors. Modelling these events separately allowed the contrast of mental-state versus sex judgments to estimate stimulus-judgment responses while accounting for the subsequent rating event. Each event was convolved with the canonical haemodynamic response function and its temporal derivative. We contrasted the mental-state judgment and sex-judgment stimulus events to define the mental-state judgment localizer regions used in the analyses.

#### Acquisition

MRI data were acquired on a Siemens MAGNETOM Vida 3T scanner (syngo MR XA60) with a 64-channel head/neck coil. A T2*-weighted gradient-echo-planar sequence with simultaneous multi-slice acceleration (multiband factor = 4) and in-plane GRAPPA acceleration (R = 2) was used. Acquisition parameters were as follows. TR = 1,160 ms, TE = 30 ms, flip angle = 63°, field of view = 210 × 210 mm, matrix = 70 × 70, 56 axial slices with 3.0 × 3.0 × 3.0 mm isotropic voxels and no inter-slice gap, partial Fourier = 7/8, phase-encoding direction anterior to posterior. A double-echo gradient-recalled echo field-mapping sequence (gre_field_mapping_2mm) was acquired in the same session to enable B0 susceptibility distortion correction (TR = 672 ms, TE1 = 4.92 ms, TE2 = 7.38 ms, flip angle = 60°, 2.0 mm isotropic voxels, matrix = 110 × 110). T1-weighted images were acquired using a three-dimensional inversion-recovery gradient-echo sequence (Siemens protocol tfl3d1_224ns; TR = 2.3 s, TE = 2.26 ms, TI = 900 ms, flip angle = 8°, 256 × 256 in-plane matrix, 192 partitions, 1-mm isotropic voxels, GRAPPA factor 2 with 24 reference lines and full Fourier acquisition). For 99 participants, the localizer and sharing task were presented as sequential epochs within one continuous BOLD acquisition; two participants required a restart and therefore contributed two runs. No volumes were manually discarded as dummy scans before first-level modelling, for which scans 1:nTR were entered. Any scanner-internal dummy acquisitions could not be determined from the available records.

### fMRI preprocessing

Functional data were preprocessed using fMRIPrep versions 25.1.4 and 25.2.4^37^, both based on Nipype 1.10.0^38^. T1-weighted images were corrected for intensity nonuniformity using N4ITK^39^, skull-stripped, segmented^40^, surface-reconstructed with FreeSurfer^41^, and normalised to MNI152NLin2009cAsym space^42,43^. Susceptibility distortion correction was performed using fMRIPrep’s fieldmap-based pipeline. The two-echo phase-difference map was unwrapped with prelude (FSL)^44^ and coregistered to the BOLD reference with antsRegistration^45^, and applied to correct distortions along the phase-encoding direction (anterior to posterior). Functional runs were corrected for susceptibility distortion, head motion corrected, coregistered to the anatomical reference by boundary-based registration^46^, and resampled to standard space in a single interpolation step. Normalised images were smoothed in SPM12 with a 6-mm full-width-at-half-maximum Gaussian kernel.

First-level nuisance regression used confound time series from fMRIPrep to account for movement-related effects^47^ and physiological noise^48^. For each run, we included 12 motion regressors comprising the six rigid-body translation and rotation estimates and their first-order temporal derivatives, six anatomical CompCor components derived from white-matter and cerebrospinal-fluid signals, and framewise displacement and DVARS as continuous regressors^49^. Volumes with framewise displacement greater than 0.5 mm, the immediately following TR and non-steady-state volumes identified by fMRIPrep were modelled using one-hot spike regressors; these volumes were not deleted. Continuous nuisance regressors were z-scored within run; spike regressors were retained as binary indicators. Participants were excluded when at least 40.0% of valid framewise-displacement samples exceeded 0.5 mm (n = 2), yielding 99 participants. A 128 s high-pass filter and SPM’s FAST autocorrelation model were applied.

### fMRI analysis

#### Analysis cohort

Of the 99 participants who met the motion criterion, seven were excluded from the fixed scenario–decision analysis cohort because of insufficient valid trials for one or both phases (n = 3), no within-participant variation in interest ratings (n = 1), or extensive spike regression that rendered the planned single-trial model non-estimable (n = 3). The final joint analyses therefore included 92 participants, each contributing 23–40 valid trials (median, 40). Within each participant, scenario and decision effects were estimated from the same set of valid trials.

#### Joint first-level model

Task effects were estimated in MATLAB R2023b using SPM12^50,51^. Scenario reading and sharing decisions were represented in the same event-related general linear model. Scenario events were modelled from scenario onset with a duration of 7 s. Decision events were modelled from decision onset using the observed response time; the response window was limited to 5 s. Recipient selection and the five post-decision rating events were entered as separate unmodulated conditions with their observed response times. When the localizer and sharing task were acquired in the same continuous run, mental-state judgment, sex-judgment and localizer-rating events were included as separate nuisance conditions.

The three-appraisal fMRI model simultaneously included interest, usefulness and emotion as first-order parametric modulators of both scenario and decision events. Each modulator was mean-centred within participant over the common scenario–decision trial set and was not standardised or residualised. Serial orthogonalisation was disabled^52^, such that each coefficient estimated the association with one appraisal dimension conditional on the other two. Because the modulators were derived from ratings obtained after each decision, they indexed BOLD activity associated with subsequently reported appraisals. All task regressors were convolved with the canonical haemodynamic response function and its temporal derivative; contrasts were defined over the canonical response. Focal contrasts quantified interest-related modulation during scenario reading, during the decision period and their direct within-participant difference (scenario minus decision). Additional appraisal and within-phase contrasts served as specificity tests.

#### Initial exploratory multiplicity

The initial age-effect screen comprised 14 models, each tested in both directions, yielding 28 directional whole-brain maps (six scenario, six decision and two recipient-selection models; Supplementary Table 12). To account for multiplicity across these maps, within-map cluster-FWE P values were additionally Bonferroni-adjusted as min(1,28*P*), equivalent to a threshold of 0.05/28 = 0.00179. The same adjustment was applied to the within-map cluster-FWE P value from the initial scenario-interest SnPM13.1.09 analysis (5,000 age-label permutations; voxel-forming P < 0.001). This was not a synchronised 28-map maximum-statistic permutation test. Joint and choice-control models were follow-up analyses in overlapping participants. Their inferential scopes were reported separately from the initial exploratory family.

#### Group-level inference

Participant-level contrast images were entered into random-effects group models. Separate models estimated the group mean, the overall cross-sectional association with age, the association with K-RMET performance and the age association conditional on K-RMET. The focal conditional model included both age and K-RMET, allowing the age coefficient to estimate age-related neural variation not shared with concurrent individual differences in social-inference performance. A motion-sensitivity model additionally included mean framewise displacement averaged across runs. Continuous covariates were mean-centred across participants. Positive and negative associations were evaluated using separate directional t-contrasts. Whole-brain maps used a voxel-forming threshold of P < 0.001 uncorrected and cluster-level family-wise error correction at P < 0.05 within each map.

#### Choice-control analysis

To determine whether interest-related modulation was retained after accounting for the eventual behavioural response, an additional first-level model included actual sharing choice (share = 1; withhold = 0) as a simultaneous parametric modulator of both scenario and decision events. Choice was mean-centred within participant over the common trial set and entered together with interest, usefulness and emotion, with serial orthogonalisation disabled. This model therefore estimated unique interest-related modulation while simultaneously accounting for the other two appraisal dimensions and eventual choice. Participant-level contrasts were evaluated using the same group-model hierarchy as the three-appraisal fMRI model.

#### Covariate and permutation-based sensitivity analyses

A targeted second-level sensitivity analysis additionally included participant sex in the model containing age, K-RMET and mean framewise displacement. This analysis was restricted to the three focal specificity contrasts: scenario interest, decision interest and scenario interest minus decision interest.

As a complementary post hoc nonparametric sensitivity analysis, the age coefficient was tested using Freedman–Lane reduced-model residual permutation. We generated 10,000 synchronised unique permutations, including the identity permutation, across the scenario-interest, decision-interest and scenario-minus-decision interest maps. The primary family compared a full model containing the intercept, age and K-RMET with a reduced model retaining the intercept and K-RMET. A separate motion-sensitivity family additionally retained mean framewise displacement in both models. Clusters were formed separately in each direction at a one-sided voxel threshold of P < 0.001 using 18-neighbour connectivity. Family-wise inference used the maximum cluster extent across all three maps and both directions; the primary and motion-sensitivity families were evaluated separately.

#### Mental-state judgment localizer analysis

The localizer was estimated in a separate event-related general linear model restricted to the localizer epoch. The first and second mental-state judgment and sex-judgment sets were modelled separately, and the valence-rating event following mental-state judgments was included as a nuisance condition. The aggregate contrast of mental-state versus sex judgments served as a task-localization and manipulation check. Because condition order was fixed, a contrast restricted to the second presentation cycle was evaluated as a sensitivity analysis. Developmental associations within the localizer were treated as secondary analyses.

#### Region-of-interest and small-volume analyses

The localizer-defined search region comprised voxels identified by the epoch-restricted group-mean contrast of mental-state versus sex judgments in 99 participants, without an age regressor. The mask included all suprathreshold voxels from clusters meeting a voxel threshold of P < 0.001 uncorrected and whole-brain cluster-level FWE P < 0.05. Within this full network, we tested the positive age association with scenario-interest modulation from the final joint model (n = 92), including K-RMET, mean framewise displacement and sex. A complementary focal analysis used an 8-mm-radius sphere centred on the group-mean localizer peak at MNI [−57, 27, 3]. These localizer-based tests used peak-level random-field-theory correction within the specified search volume^53^.

In the initial interest-only analysis (n = 97), we also evaluated the positive scenario-interest age association within the previously used 8-mm-radius sphere centred at MNI [−30, 32, −16] in left lateral orbitofrontal cortex. Meta-analytic evidence for subjective-value representations provided regional context for this anatomical target^54^; the sphere centre was not identical to the meta-analysis peak. Search-volume resels were calculated using SPM’s analytic sphere geometry or the full binary search-mask geometry and the target model’s estimated smoothness. For the orbitofrontal sphere and the initial value-union and cross-task sphere analyses reported in Supplementary Table 12, peak-FWE probabilities were obtained from SPM’s native table procedure, which returns the smaller of the random-field and voxelwise Bonferroni probabilities. All small-volume tests used positive, one-sided t-contrasts, with voxel P < 0.001 and peak-level FWE P < 0.05 within each specified volume.

#### Anatomical annotation and visualisation

Anatomical labels were assigned using the Neuromorphometrics maximum-probability atlas distributed with CAT12.9 (r2577)^55^, defined in MNI152NLin2009cAsym space at 1-mm isotropic resolution. Peak coordinates were mapped to the native atlas grid using the atlas affine and assigned the nearest voxel’s label. For cluster-level overlap, atlas labels were sampled at voxel centres of the 3-mm statistical-image grid using the image and atlas affines with nearest-neighbour interpolation. Cluster-level anatomical descriptions were distinguished from peak-voxel labels. For peaks outside grey-matter parcels, the nearest grey-matter voxel within 8 mm was identified as a proximity annotation. Neuromorphometrics provided the primary anatomical nomenclature. As a post hoc cross-check, peak locations and available significant clusters were also annotated using the CAT12-distributed AAL3 atlas^56^ in the same MNI152NLin2009cAsym space. Atlas-specific peak labels were distinguished from descriptions of cluster extent; anatomical annotation did not define the functional localizer masks or spherical regions of interest. Statistical maps were visualised using CONN 22.v2407^57^.

## Data availability

The data supporting this study will be deposited in OpenNeuro upon journal publication.

## Code availability

The analysis code supporting this study will be released on GitHub upon journal publication.

## Acknowledgements

We thank Yujin Kim for assistance with the experiments and the adolescents who participated in this study.

## Author contributions

S.J.O. and J.J. conceived and designed the study. J.L. and Y.S. contributed to the design of the behavioural and fMRI studies, respectively. S.J.O. and Y.J. performed the behavioural and fMRI experiments. S.J.O. and Y.S. performed data processing, computational analysis and statistical modelling. S.J.O. wrote the draft of the manuscript, and S.J.O., Y.S. and J.J. reviewed and edited the manuscript. J.J. supervised the study. All authors reviewed and approved the final manuscript.

