## Supplementary Information for "Perceived social value is more strongly linked to gossip decisions with age in adolescence"

Supplementary Note

Sex differences in the association between interest and sharing

Across both cohorts, an interaction between sex and interest indicated that the association between perceived interest and sharing was weaker in male than in female adolescents (Sample 1: β = −0.323, P = .02056; Sample 2: β = −0.316, P = .00665; Supplementary Table 4). The age-by-interest interaction remained positive and statistically significant in the model including sex and its interaction with interest (Supplementary Table 4).

Cross-domain structure

A one-factor confirmatory model of the eight valuation, recipient and content measures fit poorly in both cohorts (Sample 1: $\chi^{2}(22)=191.35$, CFI = .000, RMSEA = .280, SRMR = .211; Sample 2: $\chi^{2}(22)=232.93$, CFI = .039, RMSEA = .310, SRMR = .213), indicating that the specified one-factor model did not adequately describe these measures. Correlated three-factor models improved relative fit, but the Sample 1 solution was inadmissible because it contained negative residual variances (a Heywood solution), and absolute fit remained limited in Sample 2 (CFI = .779, RMSEA = .160). These models therefore do not establish three confirmed latent factors. Descriptively, the first principal component explained 30.0% and 30.6% of the variance and showed similar loading patterns across samples (absolute Tucker’s congruence coefficient |ϕ| = .981); its scores were positively associated with age in both samples ($r=.368$, $P<.001$; $r=.349$, $P<.001$). The remaining components had lower congruence coefficients across samples.

Supplementary Methods

Hierarchical Bayesian models of sharing

All models were fit with the VBA toolbox under a mixed-effects scheme (VBA-MFX), estimating participant-level parameters and group-level prior distributions. The valuation, content, and recipient-choice models followed the logistic forms given in the main-text Methods. Each model was specified separately within each sample; cross-sample analyses used only the interest, usefulness, and emotion weights common to both. For the common three-appraisal model and the models with different appraisal predictors, the prior on the group parameter mean was zero-centred with identity covariance. The VBA group-prior hyperparameters were a_vPhi = b_vPhi = 0.1 and a_sigma = b_sigma = 0.1 for each parameter.

For the valuation model, we fit models with different combinations of appraisal predictors: an intercept-only model (M0); single-rating models for interest, emotion, and usefulness (M1a–M1c); the three-rating model (M2); and the sample-specific full model adding each sample’s further ratings (M3; valence and target-effect ratings in Sample 1, and target- and recipient-perspective ratings in Sample 2). Two further models excluded the interest term: a usefulness-and-emotion model (M4) and the sample-specific full model without interest (M5). Models were compared by their free energy. Free energies for all models, and the differences attributable to the interest term (M1a − M0, M2 − M4, M3 − M5), are reported in Supplementary Table 3. The model including the sample-specific additional appraisals had the highest free energy in both samples, whereas the model including interest, usefulness and emotion was used for cross-cohort comparisons because it has the same three appraisal predictors in both cohorts.

The model including interest, usefulness and emotion was evaluated by parameter recovery and comparisons of observed and fitted sharing rates. For recovery, choices were simulated from each fitted three-appraisal model and the same model was refit to the simulated data over 100 simulations. Median recovery correlations for interest, emotion and usefulness were .916, .828 and .894 in Sample 1 and .828, .822 and .765 in Sample 2. For the comparison of observed and fitted rates, predicted sharing probabilities were compared with observed choice proportions across the 48 design cells (content $\times$ valence $\times$ target); cell-pattern correlations were .953 and .939. These checks assess rank ordering and pattern reproduction rather than exact predictive calibration; the nominal cell intervals do not incorporate parameter uncertainty. Parameter recovery and observed–fitted rate comparisons are reported in Extended Data Fig. 2.

Scale-use covariates

For each participant, three scale-use covariates were computed from the pooled set of interest, emotion, and usefulness ratings: the within-subject standard deviation of the pooled ratings, the rating range (maximum − minimum), and the proportion of responses equal to the participant’s rounded median rating. The three participant-level covariates were merged onto the trial-level dataset, z-standardised within cohort on that dataset and entered as between-subject fixed effects, with an interest $\times$ scale-use (standard-deviation) interaction, into the joint three-rating trial-level model (Supplementary Table 4).

Small-volume analysis definitions

The initial value-mask analysis used the union of seven 8-mm-radius spheres: ventromedial prefrontal cortex (MNI 0, 36, −6), deep ventromedial prefrontal cortex (0, 28, −16), medial orbitofrontal cortex (0, 50, −10), bilateral nucleus accumbens (±10, 10, −8), and bilateral lateral orbitofrontal cortex (±30, 32, −16). The saved union mask contained 539 voxels, of which 126 were within the estimated search volume of the initial scenario-interest age model (n = 97). Both the seven-sphere union and the fixed left lateral orbitofrontal sphere were evaluated in this initial interest-only age-positive map; their peak-level FWE probabilities refer to their respective search volumes.

The initial cross-task age analyses used fixed 8-mm-radius spheres centred on the age-positive peaks from the other task: the localizer-derived sphere at MNI [−39, 24, 27] was applied to the initial scenario-interest age map (n = 97), and the scenario-derived sphere at MNI [−54, 18, 39] was applied to the initial localizer age map (n = 99). These age-defined spheres are distinct from the group-mean localizer network and peak sphere used for the final joint-model analyses in Fig. 6 and Extended Data Fig. 8, which were defined without an age regressor.

For the value-union and initial cross-task analyses, peak-level FWE probabilities were obtained with SPM’s native table procedure, using analytic sphere geometry or the full saved binary-mask geometry with the target model’s estimated smoothness. The effective search-volume voxel count was the intersection with the target analysis mask. SPM’s table procedure takes the smaller of its random-field and voxelwise Bonferroni probabilities. All these tests used positive, one-sided t-contrasts, voxel P < 0.001 and peak-level FWE P < 0.05 within the specified search volume.

Additional questionnaires

In addition to the K-RMET, three self-report measures were administered to all participants in both samples: the Korean Inventory of Parent and Peer Attachment Revised (IPPA-R), the Korean Social Anxiety Scale for Adolescents (K-SAS-A), and the Korean Defining Issues Test (K-DIT).

The IPPA-R contained 75 items (25 each for mother, father and peer), rated on a 5-point Likert scale. Total attachment scores summed all 25 items for each attachment figure after reverse-scoring negatively worded items. The K-SAS-A comprises 18 items on a 5-point Likert scale, yielding Fear of Negative Evaluation, Social Avoidance and Distress in New situations, and General Social Avoidance and Distress subscales. The K-DIT presents six moral dilemmas and yields a P-score indexing postconventional moral reasoning, computed for participants meeting standard integrity criteria. Descriptive statistics and internal consistency of these measures are reported in Supplementary Table 13.

These measures did not enter the primary analyses. For a specificity comparison, each was substituted for the K-RMET in a model of the interest coefficient estimated while accounting for usefulness and emotion, using the same participants for all candidate measures within each sample. K-RMET had the lowest BIC and an estimated indirect association whose bootstrap confidence interval excluded zero in Sample 1. In Sample 2, the father-attachment score had the lowest BIC; K-RMET was ranked fifth (ΔBIC = 3.46) and its indirect interval included zero. Thus, the replicated bivariate association between K-RMET and the interest weight should be distinguished from the cohort-dependent relative ranking of candidate measures (Supplementary Table 6).

Supplementary Tables

Supplementary Table 1. Per-model sample sizes and demographics for the neuroimaging analyses

Sample size, age and sex are shown for the initial single-appraisal analyses and current joint-model and ROI analyses (Sample 2, fMRI cohort). The baseline cohort comprised 101 adolescents; two were excluded for excessive head motion. Initial-model rows refer to the main-effect analysis samples; sample sizes for other covariate specifications may differ owing to missing data. The current-analysis rows identify the samples used for Fig. 6 and Extended Data Figs. 7–8.

| **Model** | **N** | **Age, mean ± s.d.** | **Age range** | **Female** | **Male** |
| --- | --- | --- | --- | --- | --- |
| Localizer | | | | | |
| D1 mental-state judgment localizer | 99 | 14.12 ± 2.60 | 10–18 | 50 | 49 |
| D3 localizer, rating-controlled | 99 | 14.12 ± 2.60 | 10–18 | 50 | 49 |
| *Scenario phase* | | | | | |
| A1 interest | 97 | 14.19 ± 2.59 | 10–18 | 49 | 48 |
| A2 emotion | 98 | 14.15 ± 2.59 | 10–18 | 50 | 48 |
| A3 usefulness | 96 | 14.23 ± 2.56 | 10–18 | 48 | 48 |
| A4 target perspective | 98 | 14.16 ± 2.58 | 10–18 | 49 | 49 |
| A5 recipient perspective | 95 | 14.19 ± 2.57 | 10–18 | 48 | 47 |
| A6 share/withhold | 96 | 14.22 ± 2.58 | 10–18 | 49 | 47 |
| *Decision phase* | | | | | |
| B1 decision (main) | 96 | 14.22 ± 2.58 | 10–18 | 49 | 47 |
| *Recipient phase* | | | | | |
| C1 recipient (main) | 81 | 14.22 ± 2.45 | 10–18 | 40 | 41 |
| *Brain–behaviour* | | | | | |
| Localizer index × interest weight | 96 | 14.18 ± 2.60 | 10–18 | 48 | 48 |
| *Current neural analyses* | | | | | |
| Robust group-mean mental-state judgment localizer | 99 | 14.12 ± 2.60 | 10–18 | 50 | 49 |
| Joint and choice-control analyses | 92 | 14.35 ± 2.55 | 10–18 | 46 | 46 |
| Initial direct scenario-minus-decision contrast | 92 | 14.35 ± 2.55 | 10–18 | 46 | 46 |
| Matched three-phase ROI analysis (Extended Data Fig. 7a) | 79 | 14.30 ± 2.43 | 10–18 | 39 | 40 |

Behavioural Sample 1 comprised 98 adolescents (mean age 13.82 ± 2.62 years; 52 female). Whole-brain parametric maps were thresholded at voxel P < 0.001 (uncorrected) and cluster-level family-wise-error correction at P < 0.05. ROI regressions, peak-level small-volume tests and permutation analyses used the inference procedures specified in Methods.

Supplementary Table 2. Trial-level logistic models of the sharing decision

Fixed-effects estimates from logistic models of the trial-wise share/withhold choice, computed separately in Sample 1 (n = 98) and the combined-task Sample 2 behavioural subset (n = 92). Single-predictor models regressed the choice on each appraisal rating in turn. The joint model used 14,113 valid trials in Sample 1 and 11,347 valid out-of-scanner and in-scanner trials in Sample 2, excluding trials without a valid share/withhold response. It entered all three appraisal ratings (z-scored within participant), their interactions with age (z-scored within cohort on the trial-level dataset), and stimulus controls (content domain and valence as fixed-effect factors, with target included in the scenario identifier), with by-participant random intercept and interest slope and a by-scenario random intercept. Only key terms are shown; β are on the log-odds scale. Joint-model P values are from two-sided Wald z tests.

| **Model / term** | **Sample 1 (behavioural)** | | | **Sample 2 (fMRI)** | | |
| --- | --- | --- | --- | --- | --- | --- |
|  | ***β*** | ***s.e.*** | ***P*** | ***β*** | ***s.e.*** | ***P*** |
| Single-predictor models | | | | | | |
| Interest | 0.50 | 0.01 | < .001 | 0.60 | 0.01 | < .001 |
| Usefulness | 0.38 | 0.01 | < .001 | 0.50 | 0.01 | < .001 |
| Emotion | 0.06 | 0.01 | < .001 | 0.18 | 0.01 | < .001 |

Single-predictor β are per one-point rating increment; joint-model ratings are z-scored. The age × interest interaction is the trial-level counterpart of the age-related strengthening of valuation-to-decision coupling for interest. Fixed-effect stimulus-control terms (content and valence) are omitted for brevity.

#### Joint model: effect estimates and uncertainty.

P values are from two-sided Wald z tests; confidence intervals are 95% Wald intervals. Sample sizes and model specification are as above.

| Sample | Term | β | s.e. | 95% CI | z | P |
| --- | --- | --- | --- | --- | --- | --- |
| Sample 1 | Interest | 0.889 | 0.073 | [0.746, 1.032] | 12.190 | $3.49\times{10}^{-34}$ |
| Sample 1 | Usefulness | 0.450 | 0.027 | [0.397, 0.504] | 16.393 | $2.16\times{10}^{-60}$ |
| Sample 1 | Emotion | 0.125 | 0.030 | [0.065, 0.184] | 4.095 | $4.22\times{10}^{-5}$ |
| Sample 1 | Age | $-$0.0007 | 0.124 | [$-$0.243, 0.242] | $-$0.006 | 0.995 |
| Sample 1 | Age × interest | 0.377 | 0.072 | [0.235, 0.519] | 5.206 | $1.93\times{10}^{-7}$ |
| Sample 1 | Age × usefulness | 0.135 | 0.027 | [0.082, 0.189] | 4.940 | $7.80\times{10}^{-7}$ |
| Sample 1 | Age × emotion | 0.069 | 0.025 | [0.020, 0.118] | 2.758 | 0.00582 |
| Sample 2 | Interest | 0.817 | 0.063 | [0.693, 0.940] | 12.972 | $1.75\times{10}^{-38}$ |
| Sample 2 | Usefulness | 0.438 | 0.030 | [0.378, 0.498] | 14.387 | $6.27\times{10}^{-47}$ |
| Sample 2 | Emotion | 0.301 | 0.035 | [0.232, 0.370] | 8.561 | $1.11\times{10}^{-17}$ |
| Sample 2 | Age | $-$0.190 | 0.110 | [$-$0.406, 0.026] | $-$1.724 | 0.0847 |
| Sample 2 | Age × interest | 0.221 | 0.062 | [0.098, 0.343] | 3.537 | 0.000405 |
| Sample 2 | Age × usefulness | 0.074 | 0.031 | [0.014, 0.135] | 2.411 | 0.0159 |
| Sample 2 | Age × emotion | $-$0.006 | 0.027 | [$-$0.060, 0.048] | $-$0.218 | 0.828 |

Supplementary Table 3. Comparison of models with different appraisal predictors (free energy)

Free energy (a variational lower bound on the log model evidence) for each model using the same participants and trials within each sample (Sample 1, n = 98, 14,113 trials; Sample 2, n = 100, 8,400 outside-scanner trials). Higher (less negative) values indicate greater approximate model evidence. M0 contains only the intercept; M1a–M1c add a single appraisal rating; M2 adds all three; M3 adds the additional ratings specific to each sample. M4 and M5 omit the interest term. Cross-cohort analyses use the model including interest, usefulness and emotion (M2); parameter recovery and comparisons of observed and fitted sharing rates for this model are shown in Extended Data Fig. 2.

| **Model** | **Predictors** | ***Sample 1 F*** | ***Sample 2 F*** |
| --- | --- | --- | --- |
| M0 | Intercept only | −8945.69 | −5522.23 |
| M1a | Intercept + interest | −7382.28 | −4597.27 |
| M1b | Intercept + emotion | −8816.33 | −5320.99 |
| M1c | Intercept + usefulness | −7734.52 | −4842.49 |
| M2 | Intercept + interest + emotion + usefulness | −6994.76 | −4333.61 |
| M3 | Intercept + interest + emotion + usefulness + sample-specific additional ratings | −6982.18 | −4292.85 |
| M4 | Intercept + emotion + usefulness | −7662.82 | −4717.78 |
| M5 | Sample-specific full model without interest | −7611.66 | −4670.28 |

The contribution of the interest term was large in both cohorts: interest alone versus intercept (M1a − M0) ΔF = 1,563 and 925; interest beyond a usefulness-and-emotion model (M2 − M4) ΔF = 668 and 384; and interest beyond the full model without it (M3 − M5) ΔF = 629 and 377 (Sample 1 and Sample 2, respectively). The sample-specific ratings also improved fit beyond the model including interest, usefulness and emotion (M3 − M2: ΔF = 12.6 and 40.8), but they differ between cohorts and were not used for cross-sample analyses using the same three appraisal predictors.

Supplementary Table 4. Age associations with valuation-to-decision weights and robustness of the interest effect

Top: Pearson correlations between age and the participant-specific appraisal weight for each appraisal dimension in the behavioural samples (Sample 1, n = 98; Sample 2, n = 100). Bottom: the trial-level age × interest interaction (log-odds scale) in Sample 1 (n = 98) and the combined-task Sample 2 behavioural subset (n = 92), under separate robustness checks: the baseline joint model with stimulus controls, and separate extensions adding scale-use covariates, quadratic age terms, or sex and its interaction with interest. Within each sample, these models used the same valid-response trials, scenario identifiers and standardised age and appraisal variables as the primary joint model (Sample 1: 14,113 trials, n = 98; Sample 2: 11,347 trials, n = 92). Scale-use covariates were calculated from the valid-response trials. The first four rows for each sample report the age-by-interest coefficient; the final two rows report the quadratic-age-by-interest and sex-by-interest coefficients from the corresponding extended models. P values are from two-sided Wald z tests. The sensitivity models were fitted in R v4.5.2 with lme4 v1.1-37, using the primary model’s random-effects structure and optimizer settings.

| **Weight × age** | **Sample 1 (behavioural)** | | | **Sample 2 (fMRI)** | | |
| --- | --- | --- | --- | --- | --- | --- |
|  | ***r*** | ***n*** | ***P*** | ***r*** | ***n*** | ***P*** |
| Interest | .475 | 98 | < .001 | .351 | 100 | < .001 |
| Usefulness | .376 | 98 | < .001 | .169 | 100 | .094 |
| Emotion | .215 | 98 | .033 | .055 | 100 | .584 |
| Overall appraisal-weight magnitude | .517 | 98 | < .001 | .366 | 100 | < .001 |
| Relative interest weight | .311 | 98 | .002 | .181 | 100 | .071 |

| **Model or term** | $\boldsymbol{\beta}$ | **s.e.** | **95% Wald CI** | $\mathbf{z}$ | $\mathbf{P}$ |
| --- | --- | --- | --- | --- | --- |
| Sample 1 | | | | | |
| Baseline | 0.377 | 0.072 | [0.235, 0.519] | 5.206 | $<.000001$ |
| + Scale-use covariates | 0.334 | 0.073 | [0.191, 0.476] | 4.600 | .000004 |
| + Quadratic age terms | 0.381 | 0.071 | [0.242, 0.520] | 5.373 | $<.000001$ |
| + Sex and sex $\times$ interest | 0.361 | 0.071 | [0.222, 0.499] | 5.104 | $<.000001$ |
| Quadratic age $\times$ interest | $-$0.201 | 0.083 | [$-$0.364, $-$0.037] | $-$2.410 | .015974 |
| Sex $\times$ interest (male versus female) | $-$0.323 | 0.140 | [$-$0.597, $-$0.050] | $-$2.316 | .020555 |
| **Sample 2** | | | | | |
| Baseline | 0.221 | 0.062 | [0.098, 0.343] | 3.537 | .000405 |
| + Scale-use covariates | 0.215 | 0.063 | [0.090, 0.339] | 3.382 | .000721 |
| + Quadratic age terms | 0.215 | 0.064 | [0.091, 0.340] | 3.385 | .000711 |
| + Sex and sex $\times$ interest | 0.211 | 0.060 | [0.094, 0.328] | 3.528 | .000418 |
| Quadratic age $\times$ interest | $-$0.001 | 0.071 | [$-$0.139, 0.137] | $-$0.014 | .988800 |
| Sex $\times$ interest (male versus female) | $-$0.316 | 0.116 | [$-$0.543, $-$0.088] | $-$2.714 | .006650 |

For the appraisal-weight correlations with age, Fisher-z 95% confidence intervals were: interest, [0.306, 0.616] and [0.167, 0.513]; usefulness, [0.192, 0.535] and [$-$0.029, 0.353]; emotion, [0.018, 0.397] and [$-$0.143, 0.249] (Sample 1 and Sample 2, respectively). The corresponding two-sided Pearson tests have 96 and 98 degrees of freedom.

Overall appraisal-weight magnitude is the Euclidean norm of the three appraisal weights estimated jointly from interest, usefulness and emotion; relative interest weight is the interest coefficient divided by this norm. Interest weights from this three-appraisal model and overall magnitude increased with age in both cohorts. Relative interest weight showed the same positive direction in both cohorts and a significant pooled association, although the Sample 2 estimate alone was less precise. The usefulness- and emotion-weight associations were weaker and reached significance in Sample 1 only. The trial-level age × interest interaction remained significant under every robustness check (effect-size change relative to baseline: −12% and −3% with scale-use covariates; −4% and −4% in the sex-adjusted model). The quadratic-age-by-interest term was significant in Sample 1 but not in Sample 2. A sex × interest interaction was present in both cohorts (Sample 1 β = −0.323, P = .02056; Sample 2 β = −0.316, P = .00665), indicating weaker valuation-to-decision coupling for interest in male adolescents.

Supplementary Table 5. Associations between social-inference ability (K-RMET) and valuation-to-decision weights

Pearson correlations between K-RMET score and the participant-specific valuation-to-decision weight from the model including interest, usefulness and emotion for each appraisal dimension, computed separately in each cohort. Sample 2 has 98 complete K-RMET observations among the 100 participants with complete outside-scanner task data.

| **Weight** | **Sample 1 (behavioural)** | | | **Sample 2 (fMRI)** | | | **Replicated** |
| --- | --- | --- | --- | --- | --- | --- | --- |
|  | ***r*** | ***n*** | ***P*** | ***r*** | ***n*** | ***P*** |  |
| Interest | .402 | 98 | < .001 | .311 | 98 | .002 | Yes |
| Emotion | .099 | 98 | .333 | .105 | 98 | .302 | No |
| Usefulness | .216 | 98 | .033 | .145 | 98 | .154 | No |

For the K-RMET–interest-weight association, Fisher-z 95% confidence intervals were [0.221, 0.556] in Sample 1 and [0.120, 0.480] in Sample 2; both two-sided Pearson tests have 96 degrees of freedom. Replication was defined as P < 0.05 in both cohorts with consistent sign.

Supplementary Table 6. Comparison of indirect associations across candidate measures

Each candidate measure was entered between age and the interest coefficient estimated while accounting for usefulness and emotion (age → candidate → interest weight, with a direct age path). Within each cohort, all candidates used the same participant rows and outcome, making BIC comparable; ΔBIC is relative to the best-fitting candidate model. The a path is age → candidate and the b path is the candidate coefficient in the interest-weight model controlling for age. Indirect intervals and two-sided bootstrap p values were obtained from 10,000 participant-level resamples. These cross-sectional products are specificity comparisons, not causal mediation estimates. The table uses the score-available complete-case set (n = 82 and 95), including available K-DIT scores irrespective of the integrity screen. These joint complete-case sets differ from the measure-specific descriptive samples in Supplementary Table 13. Requiring a K-DIT integrity pass reduced the common comparison sets to n = 72 and 85. Indirect P values equal twice the smaller non-positive or non-negative bootstrap tail proportion, using a plus-one correction in numerator and denominator and capped at 1; they are unadjusted for comparisons across candidate measures.

| **Candidate measure** | **n** | **ΔBIC** | **a path P** | **b path P** | **Indirect result** |
| --- | --- | --- | --- | --- | --- |
| Sample 1 (behavioural cohort; common n = 82) | | | | | |
| K-RMET | 82 | 0 (best) | < .001 | .005 | β = 0.127, 95% CI [0.029, 0.260], P = 0.00860 |
| K-SAS-A | 82 | +4.91 | < .001 | .067 | β = 0.074, 95% CI [$-$0.009, 0.192], P = 0.0808 |
| IPPA-R Mother | 82 | +6.62 | .001 | .190 | β = $-$0.050, 95% CI [$-$0.134, 0.018], P = 0.134 |
| IPPA-R Father | 82 | +7.38 | < .001 | .318 | β = 0.052, 95% CI [$-$0.048, 0.165], P = 0.312 |
| K-DIT | 82 | +7.71 | .014 | .408 | β = 0.024, 95% CI [$-$0.033, 0.083], P = 0.385 |
| IPPA-R Peer | 82 | +8.37 | .877 | .821 | β = $-$0.0004, 95% CI [$-$0.0216, 0.0313], P = 0.886 |
| *Sample 2 (fMRI cohort; common n = 95)* | | | | | |
| IPPA-R Father | 95 | 0 (best) | .004 | .022 | β = 0.067, 95% CI [0.006, 0.142], P = 0.0288 |
| K-SAS-A | 95 | +2.12 | .019 | .075 | β = 0.042, 95% CI [0.0004, 0.109], P = 0.0472 |
| IPPA-R Mother | 95 | +2.32 | .229 | .084 | β = 0.021, 95% CI [$-$0.011, 0.070], P = 0.230 |
| IPPA-R Peer | 95 | +2.59 | .497 | .099 | β = $-$0.011, 95% CI [$-$0.054, 0.026], P = 0.560 |
| K-RMET | 95 | +3.46 | < .001 | .170 | β = 0.076, 95% CI [$-$0.034, 0.195], P = 0.161 |
| K-DIT | 95 | +4.87 | .102 | .471 | β = 0.012, 95% CI [$-$0.029, 0.059], P = 0.562 |

K-RMET provided the best specificity model in Sample 1 but not in Sample 2. This does not negate the significant bivariate K-RMET–interest-weight association in both cohorts (Supplementary Table 5); it shows that the unique age-adjusted association and relative ranking among candidate measures were cohort dependent.

#### **Constituent path estimates.**

Variables were standardised within each sample and analysis set. Path a is from a univariable linear regression, with df = 80 and 93 in Samples 1 and 2. Path b is from the outcome regression adjusted for age, with df = 79 and 92. The two-sided t-test P values are given in the preceding table.

| Sample | Candidate | a | s.e.(a) | t(a) | b | s.e.(b) | t(b) |
| --- | --- | --- | --- | --- | --- | --- | --- |
| Sample 1 | K-RMET | 0.410 | 0.102 | 4.019 | 0.311 | 0.106 | 2.923 |
| Sample 1 | K-SAS-A | 0.372 | 0.104 | 3.582 | 0.200 | 0.108 | 1.859 |
| Sample 1 | IPPA-R Mother | $-$0.347 | 0.105 | $-$3.313 | 0.143 | 0.108 | 1.323 |
| Sample 1 | IPPA-R Father | $-$0.452 | 0.100 | $-$4.535 | $-$0.114 | 0.114 | $-$1.004 |
| Sample 1 | K-DIT | 0.269 | 0.108 | 2.502 | 0.088 | 0.106 | 0.832 |
| Sample 1 | IPPA-R Peer | $-$0.017 | 0.112 | $-$0.155 | 0.023 | 0.102 | 0.227 |
| Sample 2 | IPPA-R Father | $-$0.296 | 0.099 | $-$2.983 | $-$0.228 | 0.098 | $-$2.322 |
| Sample 2 | K-SAS-A | 0.241 | 0.101 | 2.396 | 0.176 | 0.098 | 1.802 |
| Sample 2 | IPPA-R Mother | $-$0.125 | 0.103 | $-$1.210 | $-$0.167 | 0.096 | $-$1.745 |
| Sample 2 | IPPA-R Peer | 0.071 | 0.103 | 0.682 | $-$0.159 | 0.095 | $-$1.666 |
| Sample 2 | K-RMET | 0.497 | 0.090 | 5.529 | 0.152 | 0.110 | 1.382 |
| Sample 2 | K-DIT | 0.169 | 0.102 | 1.652 | 0.071 | 0.098 | 0.724 |

Supplementary Table 7. Robustness of the age → social inference → interest-weight indirect association across model specifications

Structural-equation models estimated the indirect association between age and the interest coefficient estimated while accounting for usefulness and emotion through K-RMET, with a direct age path, under four specifications in each cohort (n = 98 in both cohorts). All specifications included paths from age to K-RMET, from age and K-RMET to the interest weight, and from age to the emotion weight, together with residual covariance between the interest and usefulness weights. Specification D additionally included a K-RMET-to-emotion path; specification E, designated primary, further included an age-to-usefulness path; and specification F also included a K-RMET-to-usefulness path. Models used integer age and maximum-likelihood estimation. Confidence intervals are percentile bootstrap intervals from 1,000 resamples; P values are two-sided z tests based on the bootstrap standard errors. Fit indices are shown where the model is over-identified.

| **Specification** | **Cohort** | **df** | **CFI** | **RMSEA** | **SRMR** | ***Indirect β*** | **95% CI** | ***P*** |
| --- | --- | --- | --- | --- | --- | --- | --- | --- |
| A (original) | Sample 1 | 4 | .81 | .17 | .12 | 0.093 | [0.010, 0.192] | .046 |
|  | Sample 2 | 4 | .99 | .02 | .06 | 0.077 | [−0.029, 0.170] | .119 |
| D (control set) | Sample 1 | 3 | .80 | .21 | .12 | 0.094 | [0.013, 0.192] | .046 |
|  | Sample 2 | 3 | 1.00 | .02 | .06 | 0.077 | [−0.031, 0.173] | .122 |
| E (primary) | Sample 1 | 1 | 1.00 | .00 | .02 | 0.094 | [0.013, 0.192] | .046 |
|  | Sample 2 | 1 | 1.00 | .00 | .02 | 0.077 | [−0.031, 0.173] | .122 |
| F (saturated) | Sample 1 | 0 | — | — | — | 0.099 | [0.016, 0.211] | .048 |
|  | Sample 2 | 0 | — | — | — | 0.083 | [−0.026, 0.181] | .105 |

Across specifications, the indirect estimate was positive and of similar magnitude in both cohorts. In the primary specification, the interval excluded zero in Sample 1 but not Sample 2; evidence for a unique indirect association was therefore stronger in Sample 1. The bivariate age–K-RMET and K-RMET–interest-weight associations were observed in both samples. The saturated model (F) is just-identified (df = 0) and therefore has no informative fit indices.

#### **Complete estimates for the primary specification.**

Each sample included 98 participants. Coefficients refer to variables z-scored before fitting; the residual-covariance row reports a covariance, not a regression coefficient. The bivariate K-RMET–interest association and the age-adjusted K-RMET path are distinct. Confidence intervals are percentile intervals from 1,000 case-bootstrap resamples. P values are unadjusted two-sided Wald z tests using bootstrap standard errors.

| Sample | Path / parameter | Estimate | s.e. | 95% CI | z | P |
| --- | --- | --- | --- | --- | --- | --- |
| Sample 1 | Age to K-RMET | 0.385 | 0.090 | [0.209, 0.562] | 4.260 | 0.0000204 |
| Sample 1 | K-RMET to interest weight | 0.244 | 0.095 | [0.039, 0.414] | 2.576 | 0.0100 |
| Sample 1 | Age to interest weight | 0.382 | 0.085 | [0.225, 0.546] | 4.507 | 0.00000656 |
| Sample 1 | Age to emotion weight | 0.207 | 0.116 | [$-$0.026, 0.429] | 1.787 | 0.0739 |
| Sample 1 | K-RMET to emotion weight | 0.020 | 0.103 | [$-$0.173, 0.225] | 0.196 | 0.844 |
| Sample 1 | Age to usefulness weight | 0.376 | 0.079 | [0.227, 0.544] | 4.752 | 0.00000202 |
| Sample 1 | Interest–usefulness residual covariance | 0.128 | 0.074 | [$-$0.013, 0.286] | 1.732 | 0.0833 |
| Sample 1 | Indirect association | 0.094 | 0.047 | [0.013, 0.192] | 1.992 | 0.0464 |
| Sample 2 | Age to K-RMET | 0.474 | 0.095 | [0.289, 0.662] | 5.005 | 0.000000558 |
| Sample 2 | K-RMET to interest weight | 0.163 | 0.106 | [$-$0.062, 0.366] | 1.530 | 0.126 |
| Sample 2 | Age to interest weight | 0.294 | 0.102 | [0.098, 0.488] | 2.893 | 0.00382 |
| Sample 2 | Age to emotion weight | $-$0.026 | 0.123 | [$-$0.254, 0.231] | $-$0.211 | 0.833 |
| Sample 2 | K-RMET to emotion weight | 0.118 | 0.127 | [$-$0.136, 0.358] | 0.926 | 0.354 |
| Sample 2 | Age to usefulness weight | 0.160 | 0.101 | [$-$0.034, 0.362] | 1.592 | 0.111 |
| Sample 2 | Interest–usefulness residual covariance | 0.123 | 0.107 | [$-$0.087, 0.340] | 1.151 | 0.250 |
| Sample 2 | Indirect association | 0.077 | 0.050 | [$-$0.031, 0.173] | 1.547 | 0.122 |

#### **Additional model-fit statistics.**

RMSEA intervals below are 90% confidence intervals. The saturated specification has zero degrees of freedom, so its fit is not evaluated.

| Specification | Sample | $\chi^{2}$ | df | P | RMSEA [90% CI] |
| --- | --- | --- | --- | --- | --- |
| A | Sample 1 | 15.691 | 4 | 0.00346 | 0.173 [0.089, 0.266] |
| A | Sample 2 | 4.234 | 4 | 0.375 | 0.024 [0.000, 0.156] |
| D | Sample 1 | 15.660 | 3 | 0.00133 | 0.208 [0.114, 0.314] |
| D | Sample 2 | 3.174 | 3 | 0.366 | 0.024 [0.000, 0.174] |
| E (primary) | Sample 1 | 0.675 | 1 | 0.411 | 0.000 [0.000, 0.249] |
| E (primary) | Sample 2 | 0.622 | 1 | 0.430 | 0.000 [0.000, 0.245] |

Supplementary Table 8. Age associations with recipient choice

Pearson correlations between age and the participant-specific recipient-category measure for each of the five recipient categories. The hierarchical-model recipient-category intercept, estimated using all valid trials, is reported separately and has category-specific sample sizes. The proportion-based analysis uses the proportion of shared trials assigned to each category in Sample 1 (n = 98) and the complete Sample 2 outside-scanner cohort (n = 100); these proportions provide the main recipient estimates.

| **Recipient** | **Sample 1** | | | **Sample 2** | | |
| --- | --- | --- | --- | --- | --- | --- |
|  | ***r*** | ***n*** | ***P*** | ***r*** | ***n*** | ***P*** |
| Model intercepts (all valid trials) | | | | | | |
| Close friends | .24 | 81 | .033 | .27 | 98 | .007 |
| Close family | −.21 | 73 | .079 | −.27 | 85 | .012 |
| Family and friends | −.03 | 79 | .812 | −.43 | 95 | < .001 |
| Several acquaintances | −.12 | 78 | .283 | −.08 | 97 | .437 |
| Anyone (incl. strangers) | −.23 | 67 | .060 | −.32 | 90 | .002 |
| *Observed recipient proportions (sharing trials only)* | | | | | | |
| Close friends | .281 | 98 | .005 | .429 | 100 | < .001 |
| Close family | −.158 | 98 | .121 | −.256 | 100 | .010 |
| Family and friends | −.025 | 98 | .808 | −.335 | 100 | .001 |
| Several acquaintances | .003 | 98 | .973 | .180 | 100 | .074 |
| Anyone (incl. strangers) | −.187 | 98 | .066 | −.158 | 100 | .116 |

Positive r indicates a positive age association with the corresponding recipient measure; negative r indicates a negative association.

#### **Uncertainty for observed recipient proportions.**

Fisher-z 95% confidence intervals for correlations with age are shown below for the proportion-based analysis (n = 98 and 100). Two-sided Pearson tests have 96 and 98 degrees of freedom, respectively. These intervals do not apply to the separate hierarchical recipient-model rows.

| Recipient | Sample 1: 95% CI | Sample 2: 95% CI |
| --- | --- | --- |
| Close friends | [0.087, 0.454] | [0.254, 0.577] |
| Close family | [$-$0.345, 0.042] | [$-$0.431, $-$0.063] |
| Family and friends | [$-$0.222, 0.174] | [$-$0.499, $-$0.148] |
| Several acquaintances | [$-$0.195, 0.202] | [$-$0.017, 0.363] |
| Anyone, including strangers | [$-$0.371, 0.012] | [$-$0.344, 0.039] |

Supplementary Table 9. Age associations with shared content, scenario valence and targets

Pearson correlations between age and the participant-specific model weight for each content domain, scenario valence and scenario target, computed in the Sample 1 (n = 98) and Sample 2 outside-scanner (n = 100) datasets. q values are Benjamini–Hochberg false-discovery-rate (FDR) corrected across all 12 content-, valence- and target-weight tests within each cohort.

| **Content domain** | **Sample 1 (behavioural)** | | | | **Sample 2 (fMRI)** | | | |
| --- | --- | --- | --- | --- | --- | --- | --- | --- |
|  | ***r*** | ***n*** | ***P*** | ***q*** | ***r*** | ***n*** | ***P*** | ***q*** |
| Care / harm | .117 | 98 | .250 | .374 | .269 | 100 | .007 | .042 |
| Fairness / cheating | .179 | 98 | .078 | .157 | .029 | 100 | .774 | .900 |
| Loyalty / betrayal | .097 | 98 | .344 | .424 | .181 | 100 | .072 | .173 |
| Relationship | −.241 | 98 | .017 | .072 | −.058 | 100 | .569 | .900 |
| Interest / goal / achievement | −.273 | 98 | .007 | .072 | −.268 | 100 | .007 | .042 |
| Appearance | .093 | 98 | .362 | .424 | −.018 | 100 | .857 | .900 |
| Target: celebrity | .168 | 98 | .099 | .169 | −.074 | 100 | .465 | .900 |
| Target: close friend | −.088 | 98 | .389 | .424 | .029 | 100 | .772 | .900 |
| Target: disliked peer | .073 | 98 | .476 | .476 | .051 | 100 | .614 | .900 |
| Target: unfamiliar person | −.205 | 98 | .043 | .104 | −.013 | 100 | .900 | .900 |
| Negative valence | .228 | 98 | .024 | .072 | .238 | 100 | .017 | .052 |
| Positive valence | −.228 | 98 | .024 | .072 | −.238 | 100 | .017 | .052 |

Fixed-effect pooled estimates, with FDR correction across the same 12 coefficients, were r = −.270, q = .0015 for interest-, goal- and achievement-related content; r = .233, q = .0041 for negative valence; and r = .195, q = .0183 for care- and harm-related content. All other non-complementary pooled q values were at least .087. Positive and negative valence are complementary contrasts and therefore yield equal-magnitude, opposite-sign correlations.

#### **Confidence intervals for the cohort-specific correlations.**

Intervals use the Fisher-z transformation. Two-sided Pearson tests have 96 and 98 degrees of freedom in Samples 1 and 2, respectively.

| Coefficient | Sample 1: 95% CI | Sample 2: 95% CI |
| --- | --- | --- |
| Appearance | [$-$0.107, 0.286] | [$-$0.214, 0.179] |
| Care/harm | [$-$0.083, 0.309] | [0.077, 0.442] |
| Fairness/cheating | [$-$0.020, 0.364] | [$-$0.168, 0.224] |
| Interest/goal/achievement | [$-$0.447, $-$0.079] | [$-$0.441, $-$0.075] |
| Loyalty/betrayal | [$-$0.104, 0.289] | [$-$0.016, 0.364] |
| Relationship | [$-$0.420, $-$0.045] | [$-$0.251, 0.140] |
| Target: celebrity | [$-$0.032, 0.354] | [$-$0.266, 0.124] |
| Target: close friend | [$-$0.281, 0.112] | [$-$0.168, 0.224] |
| Target: disliked peer | [$-$0.127, 0.267] | [$-$0.147, 0.245] |
| Target: unfamiliar person | [$-$0.387, $-$0.006] | [$-$0.209, 0.184] |
| Negative valence | [0.031, 0.408] | [0.043, 0.415] |
| Positive valence | [$-$0.408, $-$0.031] | [$-$0.415, $-$0.043] |

The pooled correlation for interest-, goal- and achievement-related content was $-$0.270, 95% CI [$-$0.396, $-$0.135], unadjusted P = 0.000122; for negative valence, 0.233, 95% CI [0.095, 0.361], P = 0.00102; and for care- and harm-related content, 0.195, 95% CI [0.056, 0.327], P = 0.00610. Corresponding FDR-adjusted q values are reported above.

Supplementary Table 10. Age associations with recipient and content distribution indices, with minimum-trial sensitivity

Tests of Pearson correlations are two-sided. Benjamini–Hochberg correction was applied within each cohort across four recipient-distribution measures: the three entropy estimators and the ordinal recipient summary. Their adjusted q values were all 0.166 in Sample 1; in Sample 2, q = 0.0625 for each entropy estimator and q = 0.300 for the ordinal summary. No recipient-distribution association survived this within-cohort FDR correction, although all entropy estimators showed a small pooled decrease with age. The ordinal recipient-category score showed no significant pooled association with age. Content entropy and concentration changed with age only in Sample 2 and in opposite directions because concentration is the inverse of distributional dispersion; these cohort-specific estimates are not treated as replicated effects.

| **Measure** | **Sample 1 (behavioural)** | | | **Sample 2 (fMRI)** | | |
| --- | --- | --- | --- | --- | --- | --- |
|  | ***n*** | ***r* [95% CI]** | ***P*** | ***n*** | ***r* [95% CI]** | ***P*** |
| Recipient entropy (plug-in) | 98 | −.141 [−.330, .059] | .166 | 100 | −.207 [−.387, −.011] | .039 |
| Recipient entropy (Miller–Madow) | 98 | −.150 [−.338, .050] | .140 | 100 | −.202 [−.383, −.006] | .044 |
| Recipient entropy (KT shrinkage) | 98 | −.142 [−.331, .058] | .164 | 100 | −.199 [−.381, −.003] | .047 |
| Ordinal recipient-category score | 98 | −.156 [−.344, .044] | .125 | 100 | −.105 [−.295, .094] | .300 |
| Content entropy (exposure-normalised) | 98 | .010 [−.189, .208] | .925 | 100 | −.217 [−.396, −.021] | .030 |
| Content concentration (Herfindahl) | 98 | .001 [−.198, .199] | .996 | 100 | .223 [.028, .402] | .026 |

Supplementary Table 10 (continued). Minimum number of shared trials: sensitivity of recipient-measure reliability

Participants were retained only if they had at least the indicated number of shared trials. The full sensitivity analysis considered thresholds of 0, 20, 30, 40, 50, and 60 trials; 0, 30, and 60 are shown here. Reliability estimates remained similar across the displayed thresholds, although the retained sample declined. The Sample 2 rows in this supplementary threshold analysis use the paired behavioural–fMRI subset (n = 92) and are not used for the n = 100 outside-scanner main estimates.

| **Sample** | **Measure** | ***n* at 0** | ***r*_SB_ at 0** | ***n* at 30** | ***r*_SB_ at 30** | ***n* at 60** | ***r*_SB_ at 60** |
| --- | --- | --- | --- | --- | --- | --- | --- |
| Sample 1 | Recipient entropy (plug-in) | 98 | .867 | 92 | .845 | 55 | .876 |
| Sample 1 | Recipient entropy (Miller–Madow) | 98 | .858 | 92 | .836 | 55 | .874 |
| Sample 1 | Recipient entropy (KT shrinkage) | 98 | .866 | 92 | .854 | 55 | .881 |
| Sample 1 | Recipient ordinal mean | 98 | .920 | 92 | .926 | 55 | .940 |
| Sample 1 | Proportion: close friends | 98 | .899 | 92 | .881 | 55 | .891 |
| Sample 1 | Proportion: anyone | 98 | .944 | 92 | .947 | 55 | .965 |
| Sample 2 | Recipient entropy (plug-in) | 92 | .873 | 85 | .895 | 54 | .924 |
| Sample 2 | Recipient entropy (Miller–Madow) | 92 | .864 | 85 | .894 | 54 | .923 |
| Sample 2 | Recipient entropy (KT shrinkage) | 92 | .887 | 85 | .901 | 54 | .924 |
| Sample 2 | Recipient ordinal mean | 92 | .893 | 85 | .904 | 54 | .930 |
| Sample 2 | Proportion: close friends | 92 | .897 | 85 | .909 | 54 | .930 |
| Sample 2 | Proportion: anyone | 92 | .929 | 85 | .934 | 54 | .945 |

Supplementary Table 11. Cross-domain associations linking valuation weights to recipient and content indices

Within the Sample 1 (n = 98) and Sample 2 outside-scanner (n = 100) datasets, Pearson correlations were estimated between each appraisal weight from the model including interest, usefulness and emotion and each recipient/content index, before and after partialling age. Pooled estimates and 95% confidence intervals were obtained by fixed-effect Fisher-z pooling; Q tests assess between-cohort heterogeneity. P values are two-sided and are provided as a complete reference matrix rather than as a separate confirmatory family. The clearest age-adjusted cross-cohort link was between a larger interest weight and a smaller proportion of sharing to anyone.

| **Weight** | **Cross-domain measure** | **Adjustment** | ***r* Sample 1** | ***r* Sample 2** | **Pooled *r* [95% CI]** | ***P* pooled** | ***P* heterogeneity** |
| --- | --- | --- | --- | --- | --- | --- | --- |
| Interest | P(close friends) | Raw | .186 | .212 | .199 [.060, .331] | .005 | .853 |
| Interest | P(close friends) | Age-adjusted | .062 | .072 | .067 [−.075, .207] | .353 | .945 |
| Interest | P(anyone) | Raw | −.286 | −.297 | −.292 [−.415, −.158] | < .001 | .935 |
| Interest | P(anyone) | Age-adjusted | −.228 | −.261 | −.245 [−.373, −.107] | < .001 | .811 |
| Interest | Recipient ordinal mean | Raw | −.274 | −.203 | −.239 [−.367, −.101] | < .001 | .598 |
| Interest | Recipient ordinal mean | Age-adjusted | −.230 | −.178 | −.204 [−.336, −.065] | .004 | .708 |
| Interest | Recipient entropy (KT) | Raw | −.117 | −.122 | −.120 [−.256, .021] | .095 | .971 |
| Interest | Recipient entropy (KT) | Age-adjusted | −.057 | −.057 | −.057 [−.197, .085] | .430 | 1.000 |
| Interest | Content entropy | Raw | −.034 | −.211 | −.124 [−.260, .016] | .083 | .211 |
| Interest | Content entropy | Age-adjusted | −.044 | −.148 | −.096 [−.235, .045] | .182 | .469 |
| Interest | Content concentration | Raw | .045 | .217 | .133 [−.008, .268] | .064 | .224 |
| Interest | Content concentration | Age-adjusted | .051 | .152 | .102 [−.040, .240] | .157 | .481 |
| Usefulness | P(close friends) | Raw | −.025 | .078 | .027 [−.114, .167] | .708 | .472 |
| Usefulness | P(close friends) | Age-adjusted | −.148 | .007 | −.070 [−.209, .072] | .333 | .285 |
| Usefulness | P(anyone) | Raw | −.098 | −.039 | −.068 [−.207, .073] | .342 | .681 |
| Usefulness | P(anyone) | Age-adjusted | −.031 | −.013 | −.022 [−.162, .120] | .765 | .902 |
| Usefulness | Recipient ordinal mean | Raw | −.050 | .022 | −.013 [−.153, .128] | .855 | .618 |
| Usefulness | Recipient ordinal mean | Age-adjusted | .010 | .041 | .026 [−.116, .166] | .724 | .832 |
| Usefulness | Recipient entropy (KT) | Raw | .038 | .073 | .056 [−.085, .195] | .437 | .809 |
| Usefulness | Recipient entropy (KT) | Age-adjusted | .100 | .111 | .105 [−.036, .243] | .145 | .941 |
| Usefulness | Content entropy | Raw | .034 | −.204 | −.088 [−.225, .054] | .224 | .095 |
| Usefulness | Content entropy | Age-adjusted | .033 | −.174 | −.073 [−.212, .069] | .316 | .150 |
| Usefulness | Content concentration | Raw | −.024 | .208 | .094 [−.047, .232] | .190 | .102 |
| Usefulness | Content concentration | Age-adjusted | −.027 | .178 | .077 [−.065, .216] | .285 | .156 |
| Emotion | P(close friends) | Raw | .091 | .016 | .053 [−.088, .192] | .463 | .603 |
| Emotion | P(close friends) | Age-adjusted | .032 | −.009 | .011 [−.130, .152] | .875 | .778 |
| Emotion | P(anyone) | Raw | .042 | .042 | .042 [−.099, .181] | .560 | .998 |
| Emotion | P(anyone) | Age-adjusted | .085 | .052 | .068 [−.074, .208] | .345 | .815 |
| Emotion | Recipient ordinal mean | Raw | −.043 | .049 | .003 [−.137, .144] | .964 | .524 |
| Emotion | Recipient ordinal mean | Age-adjusted | −.010 | .055 | .023 [−.119, .164] | .753 | .655 |
| Emotion | Recipient entropy (KT) | Raw | −.024 | .140 | .060 [−.082, .198] | .408 | .253 |
| Emotion | Recipient entropy (KT) | Age-adjusted | .007 | .155 | .082 [−.060, .221] | .257 | .305 |
| Emotion | Content entropy | Raw | .197 | .134 | .165 [.026, .299] | .021 | .657 |
| Emotion | Content entropy | Age-adjusted | .199 | .150 | .175 [.034, .308] | .015 | .727 |
| Emotion | Content concentration | Raw | −.194 | −.122 | −.158 [−.292, −.018] | .027 | .611 |
| Emotion | Content concentration | Age-adjusted | −.199 | −.138 | −.168 [−.302, −.028] | .019 | .669 |

Supplementary Table 12. Whole-brain and small-volume neuroimaging results (Sample 2)

Whole-brain results survived a voxel-level threshold of P < 0.001 (uncorrected) and cluster-level family-wise-error (FWE) correction at P < 0.05; the P_FWE column reports cluster-level values for these rows. For the small-volume analyses, the P_FWE column reports peak-level values following the same voxel threshold. The fixed left lateral OFC sphere, seven-sphere union and initial cross-task sphere rows report native SPM-table peak-level FWE probabilities. Initial age-defined cross-task analyses and current group-mean localizer analyses with regions defined without an age regressor are reported as distinct model families. In the SVC rows, k is the contiguous suprathreshold cluster extent, not the mask size; a dash indicates that an individual peak-cluster extent is not reported.

For the current joint interest maps adjusted for age, K-RMET, mean framewise displacement and sex, T statistics have 87 residual degrees of freedom. The age-, K-RMET- and mean-framewise-displacement-adjusted joint and actual-choice-control maps have 88 residual degrees of freedom. These degrees of freedom apply to these specified model families, not to every row in this table.

The current full-network localizer mask comprises 8 clusters and 1,869 voxels (1,850 within the target analysis mask); the focal group-mean-localizer sphere is centred at MNI −57, 27, 3 with an 8-mm radius. The fixed left lateral OFC sphere was centred at MNI −30, 32, −16; MNI −33, 27, −15 is the observed peak in left posterior orbital gyrus. The sphere contained 80 nominal and 31 effective voxels, and the union mask contained 539 nominal and 126 effective voxels. For the cross-task analyses, the mental-state judgment localizer peak sphere was centred at MNI −39, 24, 27 and contained 81 nominal and effective voxels, whereas the scenario-interest peak sphere was centred at MNI −54, 18, 39 and contained 81 nominal and 50 effective voxels in the localizer age model. Sphere centres are functional ROI definitions and are distinguished from observed peaks.

Anatomical labels for the final joint analyses were assigned with the CAT12 Neuromorphometrics maximum-probability atlas in MNI152NLin2009cAsym space; the final joint-model left inferior frontal cluster (k = 116) was described using cluster-level atlas overlap. Its peak (MNI −39, 30, −9) was labelled as left cerebral white matter. Of the 116 cluster voxels, 40 (34.5%) were assigned to left cerebral white matter, 25 (21.6%) to the orbital part of the left inferior frontal gyrus, 21 (18.1%) to the left posterior orbital gyrus, 18 (15.5%) to the left frontal operculum and 6 (5.2%) to the triangular part of the left inferior frontal gyrus; the remaining 6 voxels overlapped other parcels. These percentages describe atlas assignments, not independently measured tissue composition. The actual-choice control cluster is reported separately (k = 84). Region names describe cluster extent where a matching significant-cluster export was available. For the initial localizer age-positive and age-negative contrasts and the initial scenario-minus-decision contrast at MNI [-9, 42, 30], only the peak label was verified. White-matter and ventricular assignments refer to atlas labels rather than participant-specific tissue segmentations. Neuromorphometrics supplied the primary names; AAL3 annotations provided a post hoc cross-check. Atlas labels at some peaks differed, including the full-network SVC peak labelled as frontal operculum in Neuromorphometrics and pars triangularis in AAL3. Coordinates are in MNI space. L = left; R = right.

| **Region** | **k** | **Peak T** | ***P*^FWE^** | **x** | **y** | **z** |
| --- | --- | --- | --- | --- | --- | --- |
| Current epoch-restricted localizer group mean (mental-state judgment > sex judgment), N = 99 | | | | | | |
| L inferior frontal gyrus, pars triangularis | 813 | 10.98 | < .000001 | −57 | 27 | 3 |
| L middle temporal gyrus | 465 | 8.45 | < .000001 | −63 | −48 | 6 |
| Bilateral caudate cluster with ventricular and white-matter overlap (ventricular peak) | 211 | 5.97 | < .000001 | −6 | 6 | 6 |
| R superior temporal gyrus | 109 | 6.78 | .000162 | 57 | 6 | −12 |
| R middle temporal gyrus | 91 | 6.24 | .000595 | 51 | −39 | 0 |
| L supplementary motor cortex | 76 | 5.36 | .001875 | −9 | 21 | 45 |
| Anterior cingulate cluster with predominant white-matter overlap (white-matter peak) | 54 | 5.00 | .011686 | 12 | 39 | 0 |
| R inferior frontal gyrus, pars triangularis | 50 | 6.32 | .016684 | 54 | 30 | 0 |
| *Initial full-acquisition event-related localizer (mental-state judgment > sex judgment), N = 99* | | | | | | |
| L inferior frontal gyrus, pars triangularis | 1088 | 12.23 | < .001 | −54 | 27 | 6 |
| L superior/middle temporal gyri | 499 | 10.37 | < .001 | −60 | −48 | 3 |
| R middle/superior temporal gyri | 115 | 7.93 | < .001 | 51 | −36 | 3 |
| L supplementary motor cortex extending to medial superior frontal gyrus | 94 | 7.12 | < .001 | −9 | 21 | 42 |
| R superior temporal gyrus extending to temporal pole | 74 | 6.99 | .003 | 57 | 6 | −12 |
| R cerebral white matter | 39 | 4.92 | .049 | 12 | 39 | 3 |
| *Scenario-phase interest × age (A1), N = 97* | | | | | | |
| L middle frontal gyrus (lateral PFC) | 196 | 4.46 | < .001 | −54 | 18 | 39 |
| L occipital pole | 97 | 4.35 | .018 | −18 | −99 | −3 |
| *Initial full-acquisition localizer × age, age-positive and age-negative contrasts, N = 99* | | | | | | |
| L cerebral white matter (peak label; nearest L middle frontal gyrus; age +) | 45 | 4.11 | .028 | −39 | 24 | 27 |
| L lingual gyrus (peak label; age −) | 124 | 4.38 | < .001 | 0 | −81 | 6 |
| *Initial scenario-interest × age: fixed OFC sphere and seven-sphere union, N = 97* | | | | | | |
| L posterior orbital gyrus (lateral OFC; fixed historical sphere) | 7 | 4.31 | .000618 | −33 | 27 | −15 |
| Union value mask | 7 | 4.31 | .002514 | −33 | 27 | −15 |
| *Initial-model cross-task SVC of age-positive maps in fixed age-defined spheres* | | | | | | |
| * L middle frontal gyrus (lateral PFC): localizer sphere → sharing task (N = 97) | 10 | 3.93 | .003267 | −45 | 27 | 30 |
| L middle frontal gyrus (lateral PFC): sharing-task sphere → localizer (N = 99) | 5 | 3.53 | .015700 | −51 | 21 | 33 |
| *Current group-mean-localizer SVC: final joint scenario interest × age, N = 92* | | | | | | |
| L frontal operculum (full localizer-network SVC) | — | 4.48 | .015696 | −45 | 24 | −3 |
| L inferior frontal gyrus, pars triangularis (localizer-peak sphere SVC) | 1 | 3.21 | .040448 | −54 | 21 | 0 |
| *Initial interest-only direct scenario-minus-decision contrast × age, N = 92* | | | | | | |
| L medial superior frontal gyrus (peak label) | 203 | 4.38 | .000005 | −9 | 42 | 30 |
| L posterior orbital gyrus (lateral OFC) | 60 | 4.14 | .0174 | −36 | 27 | −15 |
| L middle temporal gyrus | 57 | 4.07 | .0216 | −57 | −21 | −12 |
| *Joint scenario-interest × age, adjusted for K-RMET, mean FD and sex, N = 92* | | | | | | |
| L inferior frontal cluster | 116 | 4.73 | .000316 | −39 | 30 | −9 |
| L occipital pole | 83 | 4.40 | .002604 | −18 | −102 | 0 |
| L precuneus/posterior cingulate cluster (white-matter peak) | 76 | 3.86 | .004211 | −15 | −48 | 30 |
| R caudate | 58 | 3.96 | .015549 | 15 | 18 | 6 |
| L middle cingulate cluster with predominant white-matter overlap (white-matter peak) | 50 | 4.10 | .028883 | −3 | −18 | 24 |
| *Choice-control joint scenario-interest × age, adjusted for K-RMET and mean FD, N = 92* | | | | | | |
| L inferior frontal cluster | 84 | 4.80 | .001989 | −39 | 30 | −9 |
| Midline white matter (nearest middle cingulate gyrus) | 49 | 4.02 | .027527 | 0 | −21 | 21 |
| R frontal cluster with predominant white-matter overlap | 48 | 4.81 | .029884 | 27 | 51 | −6 |
| L occipital cluster involving occipital pole and calcarine cortex (white-matter peak) | 43 | 4.06 | .045389 | −18 | −99 | 0 |

The initial localizer model estimated event-related responses over the full acquisition; the current localizer model was restricted to the localizer epoch and separately modelled the two presentation cycles. The two initial whole-brain localizer age clusters were not retained in the age-only epoch-restricted model. In the initial-model cross-task analyses, both directions met the specified voxel and within-sphere FWE thresholds. Replacing the initial localizer target with its epoch-restricted estimate did not retain the reverse association (peak MNI [−54, 21, 33]; T = 3.11; voxel P = .001235; peak-level random-field FWE P = .057723). These cross-task age-map analyses are distinct from the group-mean localizer tests reported in Fig. 6d and Extended Data Fig. 8, for which the search regions were defined without an age regressor.

The initial interest-only direct phase contrast above predates the three-appraisal fMRI model and should not be interpreted as evidence that the age association differs between phases after accounting for usefulness and emotion. In the final joint model, the age effect on the directly constructed scenario-interest minus decision-interest coefficient had no cluster-level FWE result in the age, age-plus-K-RMET, or age-plus-K-RMET-plus-mean-FD model. A choice-control joint model yielded one age-related cluster before motion adjustment (k = 49, T = 4.64, P_FWE = .0127, MNI [21, −42, 39]) but none after adding mean FD.

As a non-parametric sensitivity analysis, we used 10,000 synchronized Freedman–Lane permutations and maximum cluster size across the three joint interest maps (scenario, decision and their difference) and both signs. No cluster survived this two-sided three-map family correction in either the age-plus-K-RMET family (smallest family-corrected P = .1152) or the age-plus-K-RMET-plus-mean-FD sensitivity family (smallest P = .1282). The corresponding parametric joint-model clusters are therefore reported as cluster-FWE findings whose robustness was examined, not as permutation-family-significant results. The initial exploratory family comprised 28 directional age-effect maps: six scenario models, six decision models and two recipient-selection models, each tested for positive and negative age associations. Scenario and decision models covered interest, emotion, usefulness, target perspective, recipient perspective and sharing choice; recipient-selection models tested the main recipient effect and intimate-versus-diffuse recipient choice. Sample sizes varied by model (n = 65–98). Only the positive scenario-interest map contained clusters passing within-map parametric cluster-FWE correction. Bonferroni adjustment across the 28 maps gave P = .0162 for the principal left middle frontal cluster (within-map P = .000577) and P = .5114 for the occipital cluster (within-map P = .018264). For the principal frontal cluster, the initial 5,000-permutation SnPM analysis gave within-map cluster-FWE P = .0028 and Bonferroni-adjusted P = .0784 across 28 maps. These Bonferroni-adjusted results are distinct from a synchronised 28-map maximum-statistic permutation analysis, which was not performed. The joint and choice-control analyses are follow-up model checks in overlapping participants, not independent replications of the exploratory finding; their within-map parametric P values and the three-map permutation results above do not constitute correction across all explored model specifications.

Supplementary Table 13. Descriptive statistics and internal consistency of additional questionnaire measures

Statistics were calculated separately within each behavioural analysis sample (Sample 1, N = 98; Sample 2, N = 100), using participants with linked questionnaire responses and complete items for the relevant scale. Sample sizes therefore vary by measure and are distinct from the joint complete-case samples used in Supplementary Table 6. IPPA-R = Inventory of Parent and Peer Attachment Revised; K-SAS-A = Korean Social Anxiety Scale for Adolescents; K-DIT = Korean Defining Issues Test; SAD = Social Avoidance and Distress.

| **Measure** | **Subscale** | **Items** | **Sample 1 (behavioural)** | | | **Sample 2 (fMRI)** | | |
| --- | --- | --- | --- | --- | --- | --- | --- | --- |
|  |  |  | ***n*** | ***α*** | ***M (SD)*** | ***n*** | ***α*** | ***M (SD)*** |
| IPPA-R | *Mother* |  |  |  |  |  |  |  |
|  | Trust | 8 | 98 | 0.923 | 31.3 (6.4) | 100 | 0.861 | 32.6 (5.3) |
|  | Communication | 7 | 98 | 0.857 | 24.3 (5.7) | 100 | 0.858 | 25.6 (5.8) |
|  | Alienation | 6 | 98 | 0.804 | 23.6 (4.3) | 100 | 0.785 | 23.9 (4.2) |
|  | Total | 25 | 98 | 0.940 | 94.1 (16.6) | 100 | 0.927 | 97.1 (15.4) |
|  | *Father* |  |  |  |  |  |  |  |
|  | Trust | 8 | 98 | 0.908 | 30.3 (6.9) | 100 | 0.929 | 31.5 (7.2) |
|  | Communication | 7 | 98 | 0.892 | 22.5 (6.7) | 100 | 0.912 | 22.0 (7.3) |
|  | Alienation | 6 | 98 | 0.836 | 22.6 (5.1) | 100 | 0.852 | 23.0 (5.1) |
|  | Total | 25 | 98 | 0.946 | 89.9 (19.1) | 100 | 0.953 | 91.5 (20.1) |
|  | *Peer* |  |  |  |  |  |  |  |
|  | Trust | 9 | 98 | 0.881 | 37.2 (5.8) | 100 | 0.906 | 36.5 (6.6) |
|  | Communication | 8 | 98 | 0.846 | 29.7 (5.9) | 100 | 0.891 | 28.8 (6.8) |
|  | Alienation | 7 | 98 | 0.703 | 26.4 (4.1) | 100 | 0.597 | 25.9 (3.9) |
|  | Total | 25 | 98 | 0.916 | 97.2 (14.2) | 100 | 0.930 | 94.8 (16.0) |
| K-SAS-A |  |  |  |  |  |  |  |  |
|  | Fear of Negative Evaluation | 8 | 87 | 0.913 | 21.7 (8.3) | 96 | 0.904 | 22.0 (7.9) |
|  | SAD (new situations) | 6 | 87 | 0.795 | 17.9 (5.3) | 96 | 0.843 | 16.8 (5.7) |
|  | SAD (general) | 4 | 87 | 0.756 | 8.7 (3.4) | 96 | 0.748 | 8.3 (3.2) |
|  | Total | 18 | 87 | 0.917 | 48.3 (14.5) | 96 | 0.908 | 47.1 (13.8) |
| K-DIT |  |  |  |  |  |  |  |  |
|  | P-score (post-conventional) | 6 dil. | 87 | — | 36.1 (18.0) | 88 | — | 36.7 (15.6) |

IPPA-R total scores include all 25 items per attachment figure after reverse scoring; subscales use their designated items. K-SAS-A total scores include 18 items. K-DIT statistics include only participants meeting all four recorded integrity criteria; Cronbach’s α is not applicable to its composite P-score.

Supplementary Table 14. Reliability of participant-level behavioural measures

Appraisal-weight rows use separate participant-level logistic fits with Cauchy priors (scale 2.5), rather than refits of the hierarchical three-appraisal model; these estimates therefore describe the reliability of the non-hierarchical estimates and are not established lower bounds for the hierarchical weights. Split-half reliability was computed over 200 spread-stratified random trial splits and Spearman–Brown corrected (rSB); brackets give the empirical 95% interval across splits. The variance-ratio reliability column reports the estimate, $\text{max}\{0,(V_{\mathrm{between}}-\overline{V}_{\mathrm{within}})/V_{\mathrm{between}}\}$, where $V_{\mathrm{between}}$ is the variance of participant point estimates and $\overline{V}_{\mathrm{within}}$ is the mean within-participant variance across 200 spread-stratified bootstrap resamples. This is not a conventional ANOVA-based ICC. Sample 1 n = 98 and Sample 2 n = 100.

| **Measure** | **Sample 1 (behavioural)** | | | **Sample 2 (fMRI)** | | |
| --- | --- | --- | --- | --- | --- | --- |
|  | ***n*** | ***r*_SB_ [95% interval]** | **Variance-ratio reliability** | ***n*** | ***r*_SB_ [95% interval]** | **Variance-ratio reliability** |
| Separate participant-level logistic estimates | | | | | | |
| Interest weight | 98 | .785 [.671, .858] | .725 | 100 | .510 [.317, .635] | .315 |
| Usefulness weight | 98 | .794 [.693, .862] | .724 | 100 | .510 [.337, .666] | .256 |
| Emotion weight | 98 | .685 [.579, .761] | .625 | 100 | .768 [.641, .838] | .678 |
| Recipient entropy (plug-in) | 98 | .869 [.815, .908] | .875 | 100 | .822 [.751, .876] | .829 |
| Recipient entropy (Miller–Madow) | 98 | .861 [.801, .904] | .872 | 100 | .807 [.729, .862] | .826 |
| Recipient entropy (KT shrinkage) | 98 | .868 [.817, .907] | .874 | 100 | .830 [.762, .879] | .838 |
| Ordinal recipient-category score | 98 | .920 [.893, .947] | .922 | 100 | .850 [.792, .892] | .838 |
| Proportion: close friends | 98 | .903 [.868, .930] | .902 | 100 | .852 [.808, .886] | .856 |
| Proportion: anyone | 98 | .943 [.921, .961] | .945 | 100 | .896 [.861, .925] | .888 |
| Content entropy (exposure-normalised) | 98 | .811 [.724, .888] | .737 | 100 | .828 [.720, .886] | .732 |
| Content entropy (KT shrinkage) | 98 | .674 [.557, .780] | .574 | 100 | .659 [.509, .758] | .533 |
| Content concentration (Herfindahl) | 98 | .807 [.706, .884] | .709 | 100 | .813 [.674, .899] | .679 |
| Content-distribution s.d. | 98 | .732 [.638, .797] | .755 | 100 | .530 [.358, .678] | .622 |
