## Extended Data for "Perceived social value is more strongly linked to gossip decisions with age in adolescence"

Extended Data Fig. 1

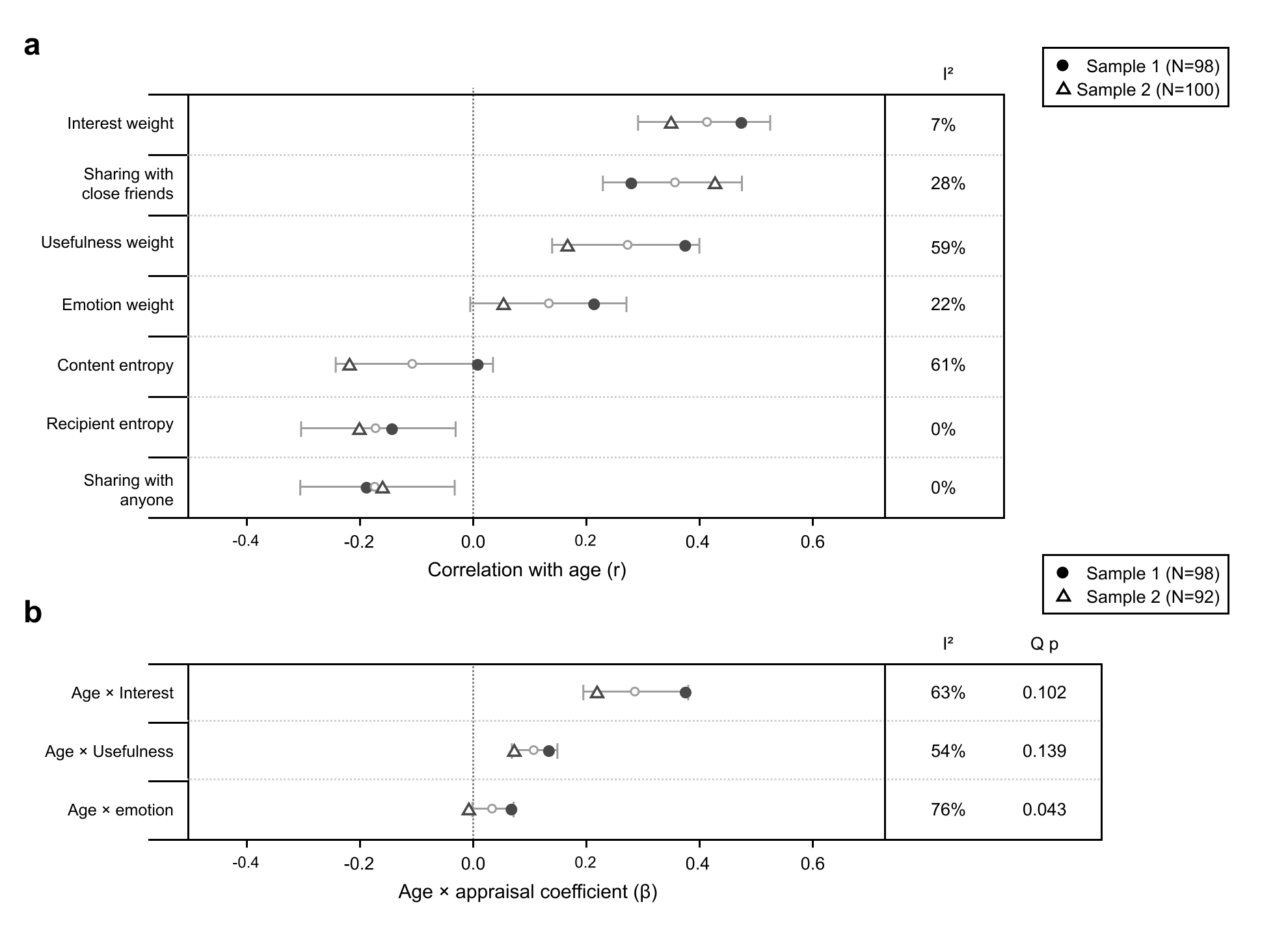

**Cross-cohort effect sizes and heterogeneity. a,** Participant-level correlations with age for common three-appraisal-model weights, conditional recipient proportions, exposure-normalised content entropy and recipient entropy calculated from Krichevsky–Trofimov-smoothed probabilities (Sample 1, n = 98; Sample 2, n = 100). Filled circles and triangles denote Sample 1 and Sample 2, respectively; small open circles and horizontal bars show fixed-effect pooled estimates and 95% confidence intervals. Interest weights were positively associated with age in both cohorts (r = 0.475 and 0.351), as was the conditional proportion of sharing directed to close friends (r = 0.281 and 0.429; pooled r = 0.358, 95% CI [0.229, 0.475], P = $2.09\times{10}^{-7}$). The usefulness-weight association was weaker in Sample 2 (r = 0.376 and 0.169), and the emotion-weight association was evident only in Sample 1 (r = 0.215 and 0.055). Sharing to anyone showed a small pooled decrease (r = −0.172, 95% CI [−0.306, −0.033], P = 0.0158), as did the displayed recipient-entropy estimator (pooled r = −0.171, 95% CI [−0.304, −0.031], P = 0.0167). **b,** Cohort-specific and pooled age-by-appraisal interaction coefficients from content- and valence-adjusted logistic mixed-effects models (Sample 1, n = 98; combined-task Sample 2 behavioural subset, n = 92). Coefficients are expressed on the log-odds scale, with age z-scored within cohort on the trial-level dataset and appraisals z-scored within participants. The age × interest interaction pooled to β = 0.287 (95% CI [0.195, 0.380], P = $1.22\times{10}^{-9}$), with no evidence that it differed between cohorts (likelihood-ratio χ²(1) = 2.43, P = 0.119). The smaller age × usefulness interaction was also positive in the pooled analysis, whereas age × emotion showed between-cohort heterogeneity (Q P = 0.043). I² and Cochran’s Q P values describe between-cohort heterogeneity. Tests of correlations and age-by-appraisal coefficients are two-sided.

Extended Data Fig. 2

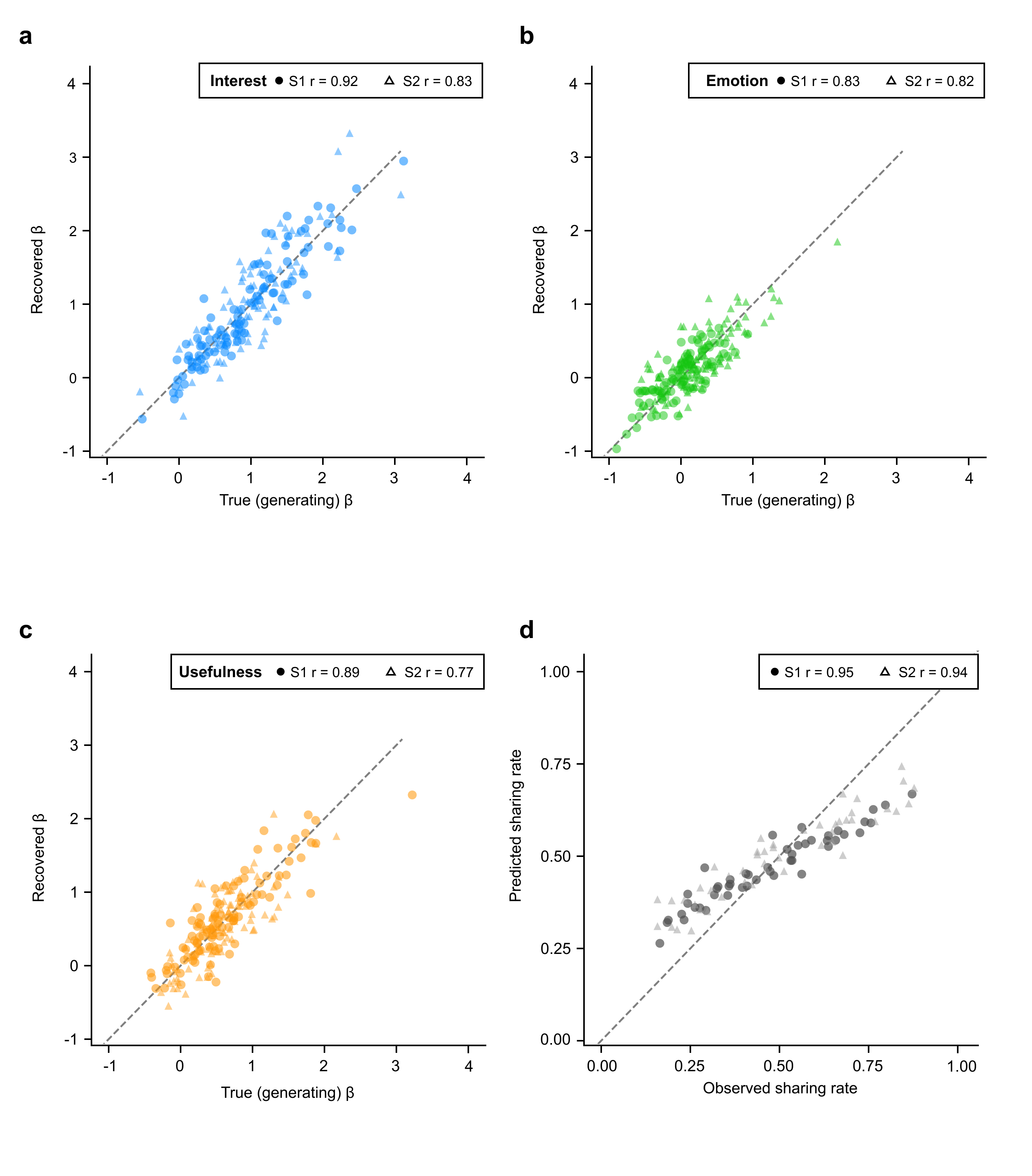

**Parameter recovery and observed versus fitted sharing rates for the three-appraisal model. a–c,** Choices were simulated from the fitted common model, containing interest (a), emotion (b) and usefulness (c), and the model was refitted in 100 simulations per cohort (Sample 1, n = 98; Sample 2, n = 100). Points show generating and recovered participant weights from simulation 1; circles denote Sample 1 and triangles denote Sample 2. The displayed r values are median Pearson correlations across all 100 simulations, rather than correlations calculated only from the displayed points: 0.916, 0.828 and 0.894 for interest, emotion and usefulness in Sample 1, and 0.828, 0.822 and 0.765 in Sample 2. **d,** Observed sharing rates versus fitted rates evaluated at posterior-mean parameters across the 48 combinations of target type, content domain and valence in each sample (Sample 1 r = 0.953; Sample 2 r = 0.939). Points represent design cells. Dashed lines in all panels indicate identity.

Extended Data Fig. 3

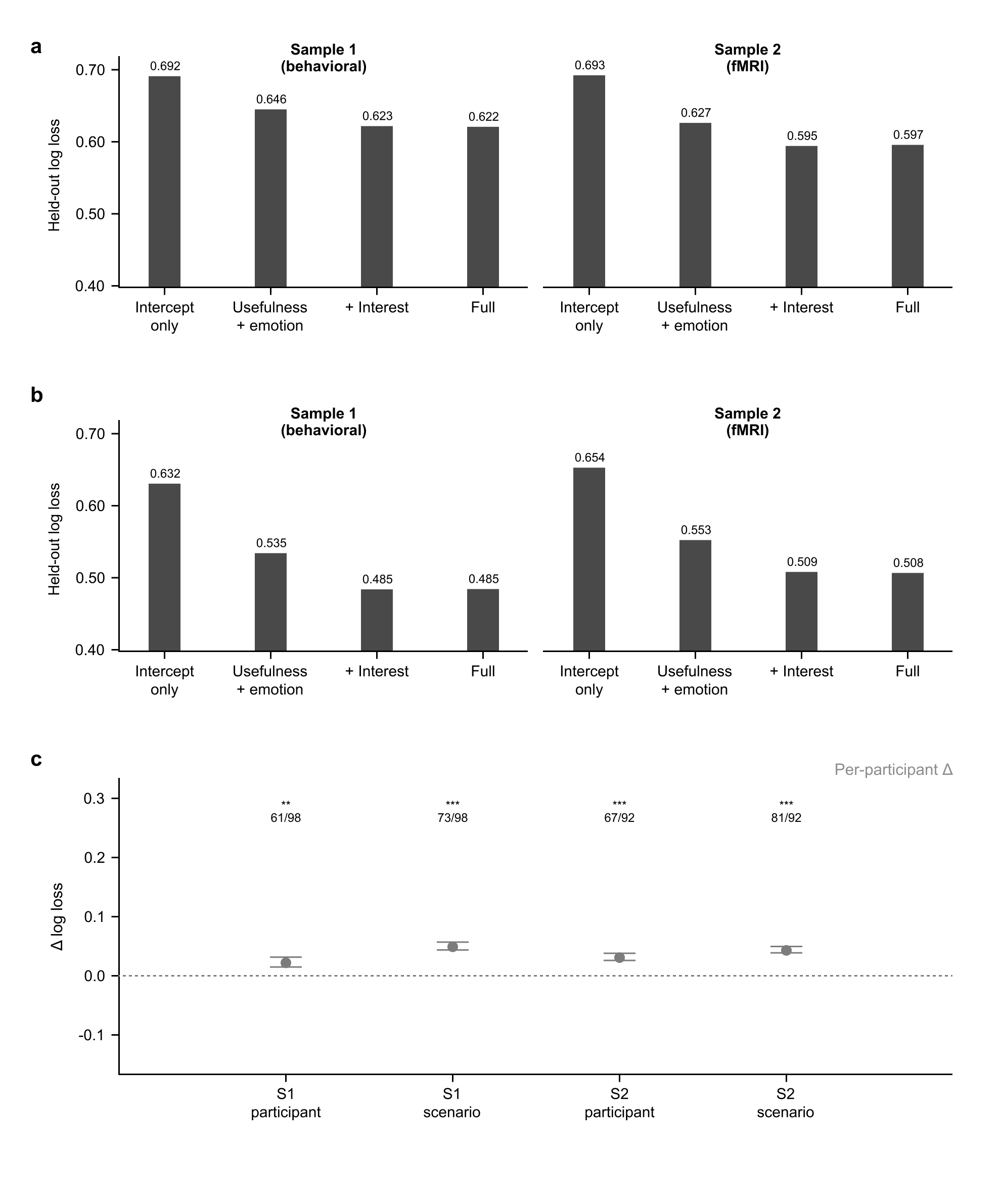

**Out-of-sample prediction of sharing choices. a and b, Ten-fold cross-validation holding out participants (a) or** combinations of target type, content domain and valence (b) (Sample 1, n = 98; combined-task Sample 2 behavioural subset, n = 92). Bars show trial-weighted held-out log loss. The models shown, in order, include an intercept only; usefulness and emotion; interest, usefulness and emotion; and these three appraisals plus cohort-specific ratings. Lower values indicate better prediction. **c,** Mean participant-specific improvement in held-out log loss after adding interest to usefulness and emotion, calculated as log loss without interest minus log loss with interest; positive values favour inclusion of interest. Points and bars show the mean $\pm$ s.e.m., and fractions give the number of participants with positive differences. Interest improved prediction in all four cohort-by-cross-validation conditions (Sample 1 participant: mean $\Delta$log loss = 0.023, 95% CI [0.0066, 0.0398], t(97) = 2.773, P = 0.0067; Sample 1 design-cell: $\Delta$ = 0.050, 95% CI [0.0372, 0.0635], t(97) = 7.591, P = $1.96\times{10}^{-11}$; Sample 2 participant: $\Delta$ = 0.032, 95% CI [0.0200, 0.0441], t(91) = 5.290, P = $8.39\times{10}^{-7}$; Sample 2 design-cell: $\Delta$ = 0.044, 95% CI [0.0335, 0.0549], t(91) = 8.227, P = $1.30\times{10}^{-12}$). **P < 0.01; ***P < 0.001, two-sided paired t-tests across participants, with 95% t-based confidence intervals for the mean paired difference.

Extended Data Fig. 4

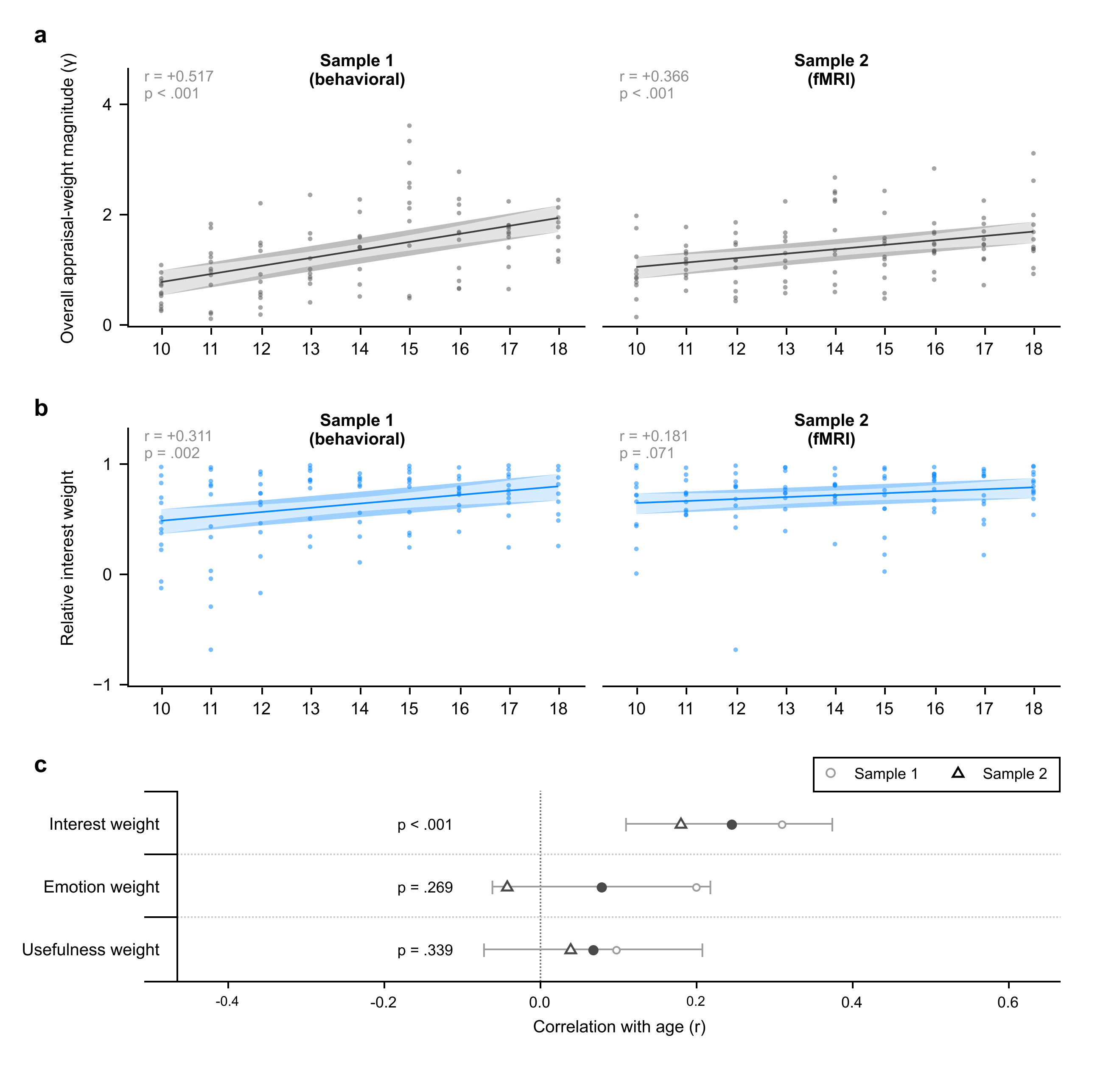

**Overall appraisal-weight magnitude and relative interest weight. a,** The Euclidean norm of each participant’s common three-appraisal weight vector ($\gamma$), excluding the intercept, was positively associated with age in both cohorts (Sample 1: n = 98, r = 0.517; Sample 2: n = 100, r = 0.366; pooled r = 0.444, 95% CI [0.324, 0.550], P = $3.76\times{10}^{-11}$). **b,** Relative interest weight, the interest coefficient divided by $\gamma$, showed a positive age association in Sample 1 (r = 0.311, P = 0.00185) and a weaker association in the same direction in Sample 2 (r = 0.181, P = 0.0714); the pooled association was r = 0.246 (95% CI [0.110, 0.374], P = 0.000494). Points represent participants. Grey and blue lines show ordinary least-squares fits for overall magnitude and relative interest, respectively; shaded bands show pointwise 95% confidence intervals for the fitted mean. **c,** Age correlations for relative interest, emotion and usefulness weights. Open circles and triangles denote Samples 1 and 2; filled circles and horizontal bars show pooled estimates and 95% confidence intervals. Pooling used inverse-variance fixed effects on Fisher-z correlations, with weights n $-$ 3. Displayed P values are unadjusted, two-sided tests of the pooled associations.

Extended Data Fig. 5

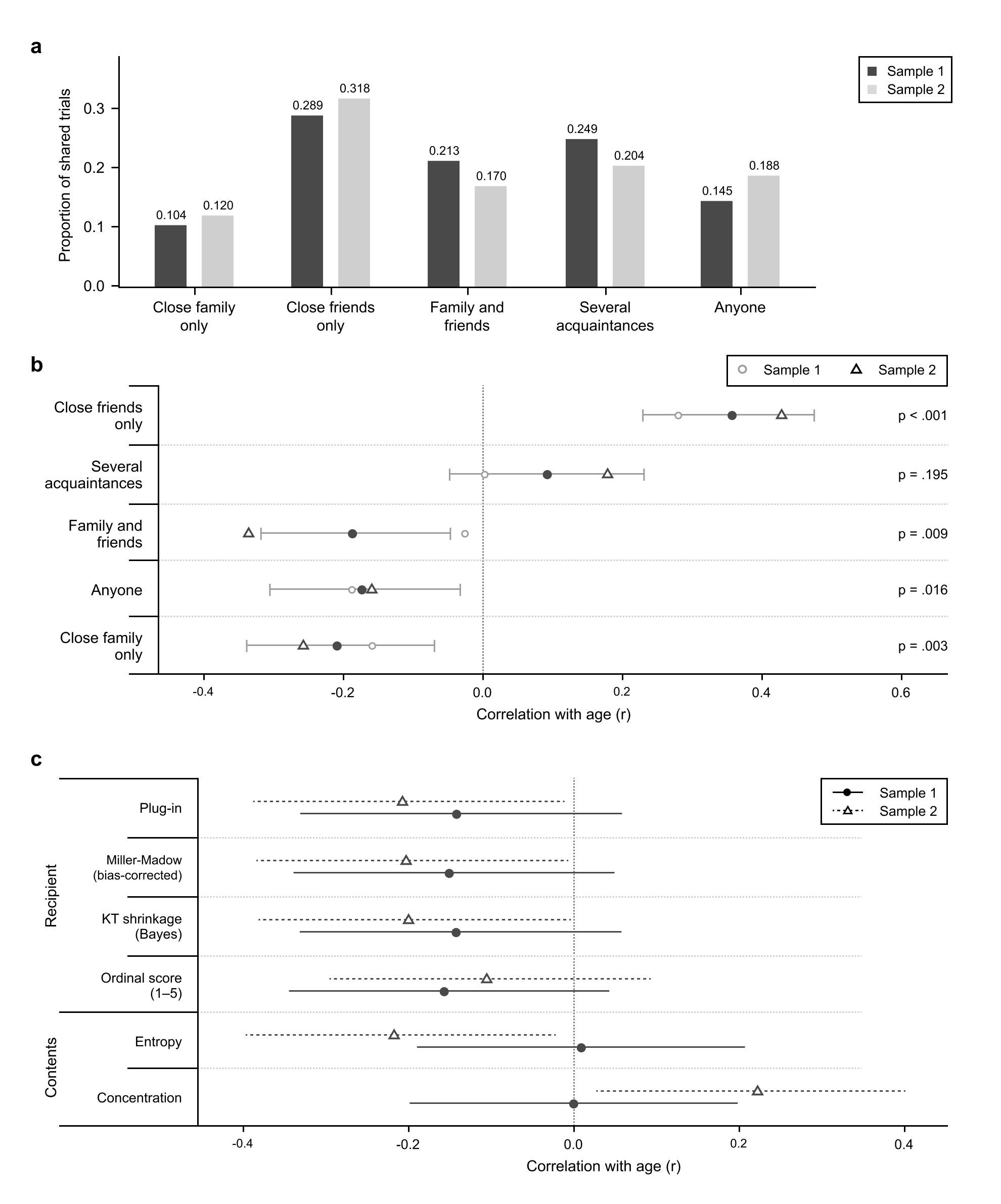

**Age associations with gossip recipient selection. a,** Mean participant-specific proportions of shared trials assigned to five recipient categories (Sample 1, n = 98, dark grey; Sample 2, n = 100, light grey). The denominator is each participant’s total number of trials on which they chose to share. **b,** Age correlations for recipient proportions. Open circles and triangles denote Samples 1 and 2; filled circles and horizontal bars show fixed-effect pooled estimates and Fisher-z 95% confidence intervals. Displayed P values are unadjusted and two-sided. The proportion of sharing choices directed to close friends was positively associated with age in both samples (r = 0.281 and 0.429; pooled r = 0.358, 95% CI [0.229, 0.475], P = $2.09\times{10}^{-7}$). Negative pooled age associations were observed for close family (r = $-$0.208, P = 0.00343), family and friends (r = $-$0.186, P = 0.00909) and anyone including strangers (r = $-$0.172, P = 0.0158); the family-and-friends association differed between cohorts (heterogeneity P = 0.0250). **c,** Age correlations for three recipient-entropy estimators, an ordinal recipient-category score, and exposure-normalised content entropy and concentration. Filled circles with solid intervals denote Sample 1; open triangles with dotted intervals denote Sample 2. Intervals are sample-specific Fisher-z 95% confidence intervals. Recipient entropy uses observed proportions (plug-in), a finite-sample bias correction (Miller–Madow), or probabilities smoothed with a pseudocount of 0.5 per category (KT shrinkage). The ordinal recipient-category score is the mean of the category codes, from 1 (close family) to 5 (anyone including strangers), across sharing trials. Content summaries use within-category sharing rates normalised across the six content categories. Pooled recipient-entropy correlations ranged from $-$0.171 to $-$0.177; the pooled ordinal association was $-$0.130 (P = 0.0695). Pooling used inverse-variance fixed effects on the Fisher-z scale, with weights n $-$ 3.

Extended Data Fig. 6

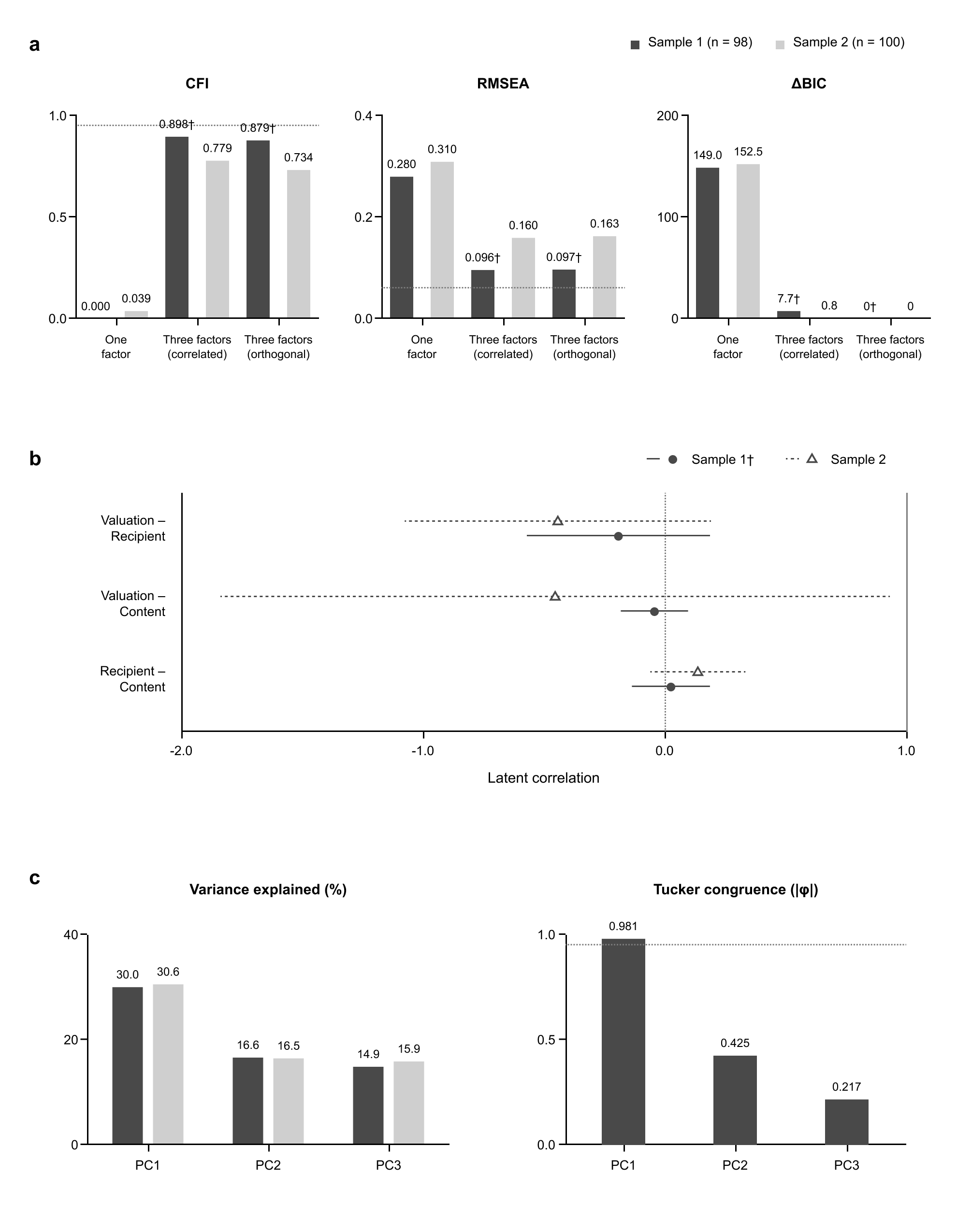

**A one-factor model fits the cross-domain measures poorly in both samples.** Analyses used eight indicators standardised within each sample (Sample 1, n = 98; Sample 2, n = 100): three common-model appraisal coefficients, four recipient measures and exposure-normalised content entropy. **a,** Comparative fit index (CFI), root mean square error of approximation (RMSEA) and relative Bayesian information criterion ($\Delta$BIC) for one-factor, correlated three-factor and orthogonal three-factor models. Models were estimated using robust maximum likelihood; displayed CFI and RMSEA are the conventional fit indices. The one-factor model fit poorly in both cohorts (CFI = 0.000 and 0.039; RMSEA = 0.280 and 0.310). Dark- and light-grey bars denote Samples 1 and 2. $\Delta$BIC is relative to the lowest BIC among the three models within each sample. Horizontal references mark CFI = 0.95 and RMSEA = 0.06. $\dagger$ denotes inadmissible Sample 1 three-factor solutions containing negative residual variances. The single-indicator content factor had its loading fixed to one and residual variance fixed to zero. **b,** Latent correlations from the correlated three-factor model. Sample 1 estimates derive from the inadmissible solution identified in a and are shown descriptively. Filled circles with solid intervals denote Sample 1 and open triangles with dotted intervals denote Sample 2. Horizontal bars are unbounded Wald 95% confidence intervals, without truncation to [$-$1, 1]. **c,** Variance explained by the first three principal components and absolute Tucker’s congruence coefficients ($|\phi|$) comparing component loading patterns across samples. PC1 explained 30.0% and 30.6% of the variance, with $|\phi|$ = 0.981; PC2 and PC3 had $|\phi|$ = 0.425 and 0.217. The horizontal reference marks $|\phi|$ = 0.95.

Extended Data Fig. 7

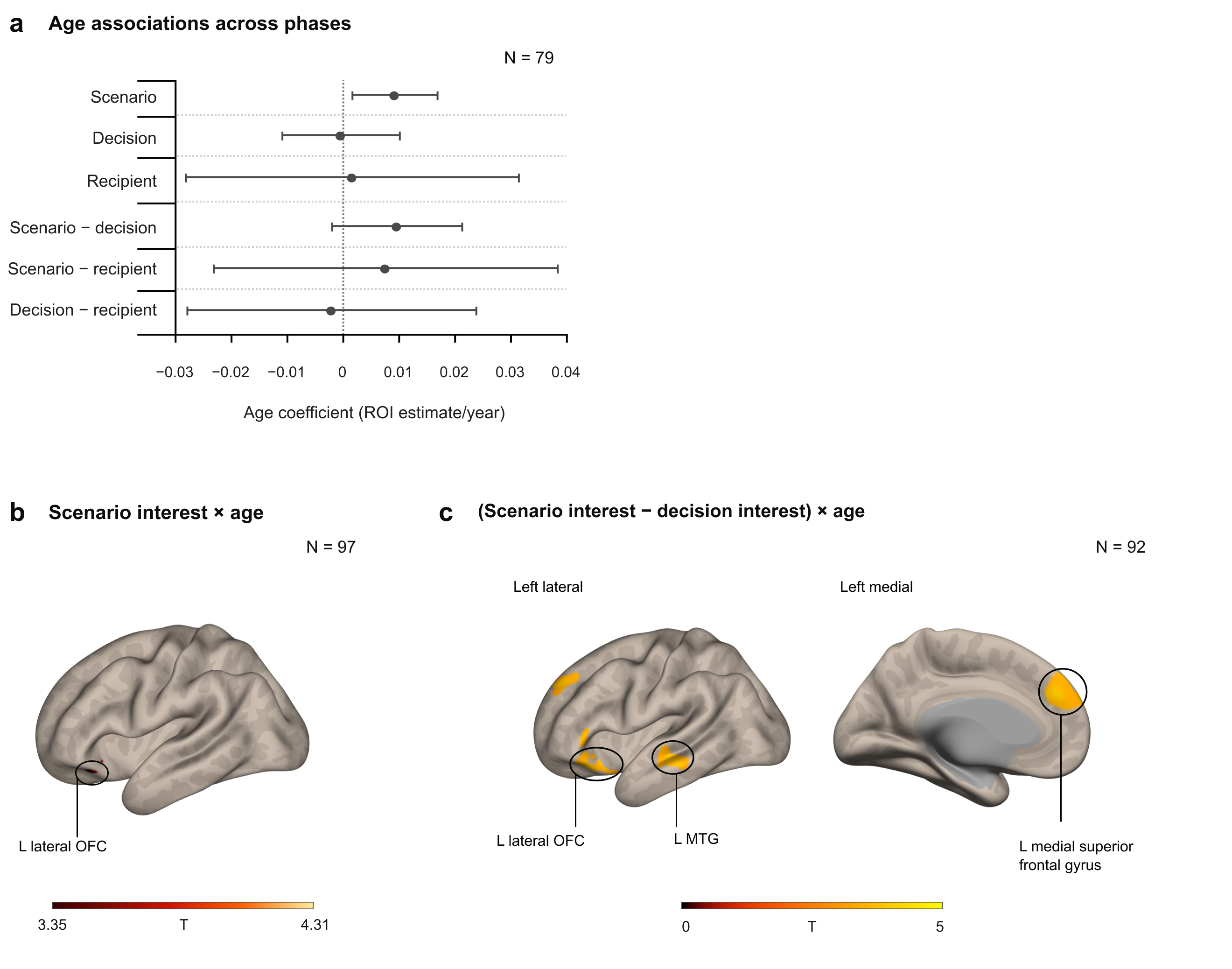

**Phase sensitivity and value-related age associations. a,** Same-participant phase comparisons in an 8-mm sphere centred on a peak from the initial localizer age-association analysis (MNI [$-$39, 24, 27]; n = 79). Age terms for scenario reading, sharing decisions and recipient selection and all three direct phase contrasts were estimated in the same participants with sex and mean framewise displacement included as covariates. Points are adjusted age coefficients in ROI-estimate units per year; bars are 95% confidence intervals; the vertical dotted line marks zero. The scenario age term was positive (P = 0.0179), whereas all direct phase contrasts were P $\geq$ 0.103. This historical age-defined ROI is distinct from the group-mean localizer sphere used for the Fig. 6 participant plots, whose centre was selected without an age regressor. **b,** Value-mask small-volume correction for scenario interest $\times$ age in the initial interest-only model (n = 97). The fixed 8-mm left lateral orbitofrontal sphere from the previously used seven-sphere value mask was centred at MNI [$-$30, 32, $-$16]; the observed peak was in left posterior orbital gyrus (MNI [$-$33, 27, $-$15]; k = 7; T = 4.31; native SPM peak-level P_FWE-SVC_ = 0.00062). **c,** Whole-brain direct (scenario interest $-$ decision interest) $\times$ age contrast from the initial interest-only model (n = 92), thresholded at voxel P < 0.001 uncorrected and cluster-level FWE P < 0.05. Clusters were observed in left medial superior frontal gyrus (MNI [$-$9, 42, 30]; k = 203; T = 4.38; P_FWE_ = 0.000005), left posterior orbital gyrus (MNI [$-$36, 27, $-$15]; k = 60; T = 4.14; P_FWE_ = 0.0174) and left middle temporal gyrus (MNI [$-$57, $-$21, $-$12]; k = 57; T = 4.07; P_FWE_ = 0.0216). This initial-model phase contrast was not reproduced in the joint appraisal model. The separate synchronized 10,000-permutation analysis evaluated the three joint-model interest maps across both directions; no cluster survived family-wise correction. Panel c shows left lateral and medial views. Black outlines locate approximate surface regions, not inferential boundaries. Colour scales indicate surface-display T statistics (b, 3.35–4.31; c, 0–5); zero is the colour-scale origin, not the inclusion threshold. Panel b reports peak-level FWE small-volume correction within a fixed sphere, whereas c reports whole-brain cluster-level FWE correction.

Extended Data Fig. 8

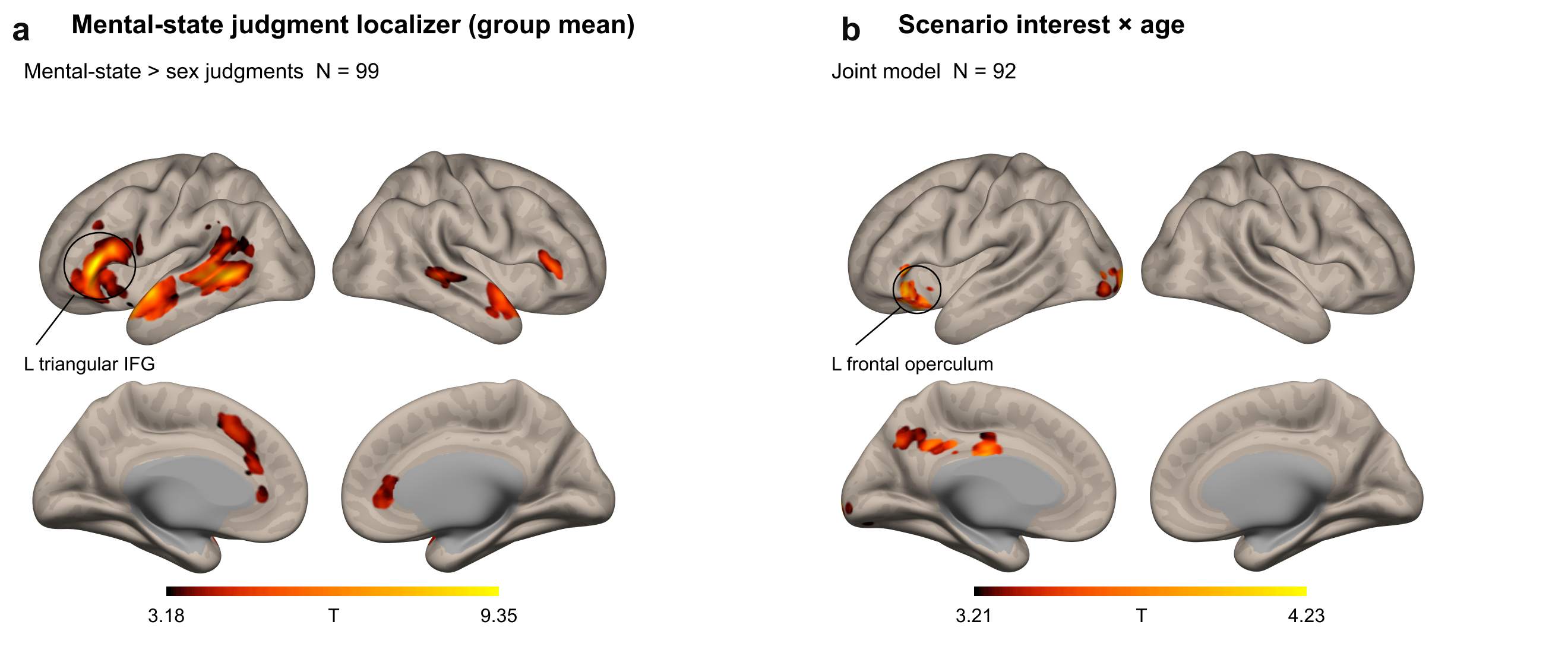

**Age-related interest modulation is observed within regions defined by the mental-state judgment localizer. a,** Surface views of the epoch-restricted group-mean contrast of mental-state versus sex judgments in a separate task (n = 99). The mental-state judgment localizer was defined without an age regressor in the same cohort. The localizer peak lies in left inferior frontal gyrus (pars triangularis) (MNI [$-$57, 27, 3]). **b,** Surface views of the three-appraisal scenario-interest age-association map (n = 92). Interest, usefulness and emotion were modelled simultaneously without serial orthogonalisation at first level; the group model included age, K-RMET, mean framewise displacement and sex. Within the full significant group-mean localizer network, the age-positive association was detected in left frontal operculum (MNI [$-$45, 24, $-$3]; T = 4.48; peak-level P_FWE-SVC_ = 0.0157). Small-volume correction used the entire significant localizer network rather than a peak-centred sphere. Both panels provide whole-brain context at voxel P < 0.001 uncorrected and whole-brain cluster-level FWE P < 0.05; the SVC P value refers to the reported peak tested within the localizer mask. Circles and leader lines indicate approximate surface regions; MNI coordinates identify statistical peaks. Colour bars show surface-display T values (a, 3.18–9.35; b, 3.21–4.23). Anatomical descriptions use the CAT12 Neuromorphometrics atlas in MNI152NLin2009cAsym space.
